# Developmental remodeling uncouples neural stem cell function from niche anatomy

**DOI:** 10.64898/2026.07.13.738185

**Authors:** I. Mateos-White, A. Marín-Garnes, L. Veintimilla-Escot, J. Fabra-Beser, L. Lázaro-Carot, J. Planells, C.M. Mateos-Martínez, M.C. Martínez-Bisbal, E. Martínez-Martínez, S.R. Ferrón, C. Gil-Sanz

**Affiliations:** Instituto de Biotecnología y Biomedicina (BIOTECMED), Universitat de València, Burjassot, Valencia, Spain; Departamento de Biología Celular y Biología Funcional. Universitat de València, Burjassot, Valencia, Spain; Centro de Investigación Biomédica en Red sobre Enfermedades Neurodegenerativas (CIBERNED), Universitat de València, Burjassot, Valencia, Spain; Instituto Interuniversitario de Investigación de Reconocimiento Molecular y Desarrollo Tecnológico (IDM), Universitat Politècnica de València - Universitat de València, Valencia, Spain; Departamento de Química Física, Universitat de València, Valencia, Spain; Unidad Mixta de Investigación Cerebrovascular, Universitat de València - Instituto de Investigación Sanitaria La Fe de Valencia, Valencia, Spain; CIBER de Bioingeniería, Biomateriales y Nanomedicina, Instituto de Salud Carlos III, Spain

## Abstract

Stem cells are generally thought to depend on specialized niches that provide signals for their long-term maintenance. Whether stem-cell competence is intrinsically constrained by anatomical organization remains unclear. Here, we show that neural stem cells can establish and sustain functional persistent stem-cell populations outside their normal anatomical context. Using developmental deletion of the cell-adhesion regulator *Afadin* as a tool to disrupt cortical tissue organization, we find that neural progenitors are displaced from the ventricular surface and establish an ectopic germinal zone (EGZ) into advanced adulthood. EGZ stem-cell populations retain self-renewal and multilineage differentiation capacity throughout this period. In parallel, the ventricular-subventricular zone (V-SVZ) undergoes persistent reorganization of tissue architecture, cellular composition, molecular state, and stem-cell activity. Together, these findings reveal unexpected plasticity in the relationship between stem cells and their tissue environment, suggesting that canonical niche anatomy may constrain where stem cells normally reside without defining the limits of functional stem-cell competence.

## Introduction

Stem cell function depends not only on intrinsic cellular programs but also on specialized microenvironments in which stem cells reside. Across tissues, stem cell niches are traditionally viewed as anatomically defined environments that regulate stem cell maintenance, self-renewal, differentiation, and tissue homeostasis throughout life^1–3^. Whether these niches are permanently established during development or retain the capacity to adopt alternative functional states after developmental perturbations remains a fundamental unanswered question in stem cell biology.

Adult neurogenesis persists in the mammalian brain through the activity of a small and highly regulated population of neural stem cells (NSCs) maintained within specialized regions, including the ventricular-subventricular zone (V-SVZ) lining the lateral ventricles and the subgranular zone of the hippocampal dentate gyrus^4–6^. Within these environments, NSCs are maintained in a dynamic balance between quiescence and activation, ensuring lifelong plasticity while preventing stem cell exhaustion or aberrant growth^7^. This balance emerges from the integration of intrinsic transcriptional programs with extrinsic signals provided by cerebrospinal fluid (CSF), vasculature, extracellular matrix, and neighboring cells^8–10^. Therefore, the functional identity of a stem cell niche may reflect not only its molecular composition but also the developmental context in which cellular interactions are established. How these diverse cues are structurally integrated to preserve NSC identity and long-term homeostasis remains a central question in adult stem cell biology. In particular, it remains unknown whether the canonical anatomical organization of neurogenic niches represents an absolute requirement for stem cell function or a developmental configuration that can be remodeled while preserving stem-cell competence^5,11,12^.

In the V-SVZ, NSCs retain apical contact with the ventricular surface and basal interactions with blood vessels, forming a polarized architecture that is essential for niche integrity^13,14^. Coordinated interactions between NSCs, neighboring cells, and their extracellular microenvironment provide structural and signaling cues that regulate NSC behavior^15–17^. Although both cell-matrix and cell-cell interactions contribute to niche organization, here we focus on cell-cell adhesion because of its central role in establishing tissue architecture and polarity. Apical adherens junctions between ependymal cells and NSCs assemble into characteristic pinwheel structures that anchor NSCs to the ventricular surface and contribute to quiescence regulation^13^. For example, *Vcam1* anchors NSCs to the apical niche and promotes quiescence through redox signaling^18^, while *Cdh2* maintains apico-basal polarity and restrains proliferation^19^. Disruption of these cell-cell adhesion systems destabilizes niche architecture, highlighting the close relationship between tissue organization and NSC regulation.

Adult V-SVZ NSCs originate from early specified embryonic radial glial cells (RGCs) that persist into adulthood as quiescent progenitors^20^. Notably, many adhesion molecules essential for adult NSC regulation are already operative in embryonic RGCs. *Vcam*1 loss reduces progenitor maintenance and accelerates neuronal differentiation during development^21^. Similarly, *Cdh2* regulates RGC proliferation and polarity in cooperation with cytoplasmic adaptor protein AFADIN^22^. *Afadin* (*Mllt4/Af6*), a core component of adherens junctions, links cadherins to the actin cytoskeleton and integrates adhesive cues with intracellular signaling^23–25^. Early loss of *Afadin* in the dorsal telencephalon disrupts RGC adhesion, leading to cortical hyperplasia, defective neuronal migration, and double-cortex formation^22^, a malformation also observed in human neurodevelopmental disorders^26^. Despite its established developmental role, whether *Afadin*-dependent regulation of tissue architecture influences the long-term organization and function of adult stem cell niches has remained unknown. Given the developmental continuity between RGCs and adult NSCs, and the importance of adherens junctions in organizing the V-SVZ niche, *Afadin* provides a unique opportunity to test how early tissue organization shapes the long-term architecture and function of stem cell environments.

Here, we use developmental loss of *Afadin* as an experimental model to test whether canonical anatomical niches strictly constrain neural stem cell function or whether stem cells retain the capacity to maintain stem-cell properties within alternative tissue environments. Using an embryonic conditional deletion approach, we show that developmental loss of *Afadin* durably remodels adult neural stem cell niches and reveals unexpected plasticity in their organization and regulation. Early dorsal loss of *Afadin* redirects a subset of neural progenitors into a persistent ectopic germinal zone (EGZ) in the cortex, outside the ventricular surface, while simultaneously remodeling the endogenous V-SVZ. These changes are accompanied by regional alterations in ventricular and ependymal organization, stem-cell activation, and gene expression. Mosaic deletion of *Afadin* in postnatal and adult NSCs further demonstrates that *Afadin* also contributes intrinsically to the regulation of NSC activation within an otherwise established niche. Together, our findings reveal that developmental perturbations can durably rewrite the spatial organization of adult neural stem cell niches, while neural stem cells retain the intrinsic competence to sustain stem-cell function beyond their canonical anatomical context.

## Results

### Developmental cortical disorganization establishes an ectopic neurogenic niche in the adult brain

To determine whether developmental disruption of tissue organization can durably alter the spatial organization of neural stem cell (NSC) niche, we examined the consequences of embryonic *Afadin* deletion in adult mice. We previously showed that conditional inactivation *of Afadin* in the dorsal telencephalon in *Afadin^Flox/Flox^; Emx1^CRE/+^* (*Afadin^cKO^)* causes RGC overproliferation during neocortical development^22^. Because embryonic RGCs give rise to adult NSCs, we first assessed whether this developmental phenotype extends to the adult V-SVZ (Fig. 1A). Immunohistochemical analysis of the proliferation marker KI67^27^ revealed a marked increase in proliferating cells together with a profound alteration in their spatial distribution in adult *Afadin^cKO^* compared with *Afadin^Flox/Flox^* (*Afadin^Ctl^)* controls (Fig. 1B). This increase was detected in dorsal regions, where *Afadin* was disrupted (Extended Data Fig. 1A) and the V-SVZ appeared markedly expanded relative to controls (Fig. 1B, 1–1′; Extended Data Fig. 1A). Unexpectedly, elevated proliferation was also observed in ventral regions, where *Afadin* expression was preserved (Fig. 1B, 2–2′; Extended Data Fig. 1A), revealing non–cell-autonomous consequences of *Afadin* loss. Most strikingly, numerous KI67⁺ cells were found outside the V-SVZ within a structure absent from control brains: an ectopic white matter (EWM) located between the normocortex and the heterotopic cortex, far from the ventricular surface (Fig. 1B, 3–3′; Extended Data Fig. 1B, arrowheads). These proliferating cells were confined to the lateralmost EWM overlying the striatum and extended along the rostrocaudal axis (Extended Data Fig. 1B; Fig. 1C), defining a previously unrecognized proliferative domain that we hereafter term the ectopic germinal zone (EGZ).

**Figure 1.**
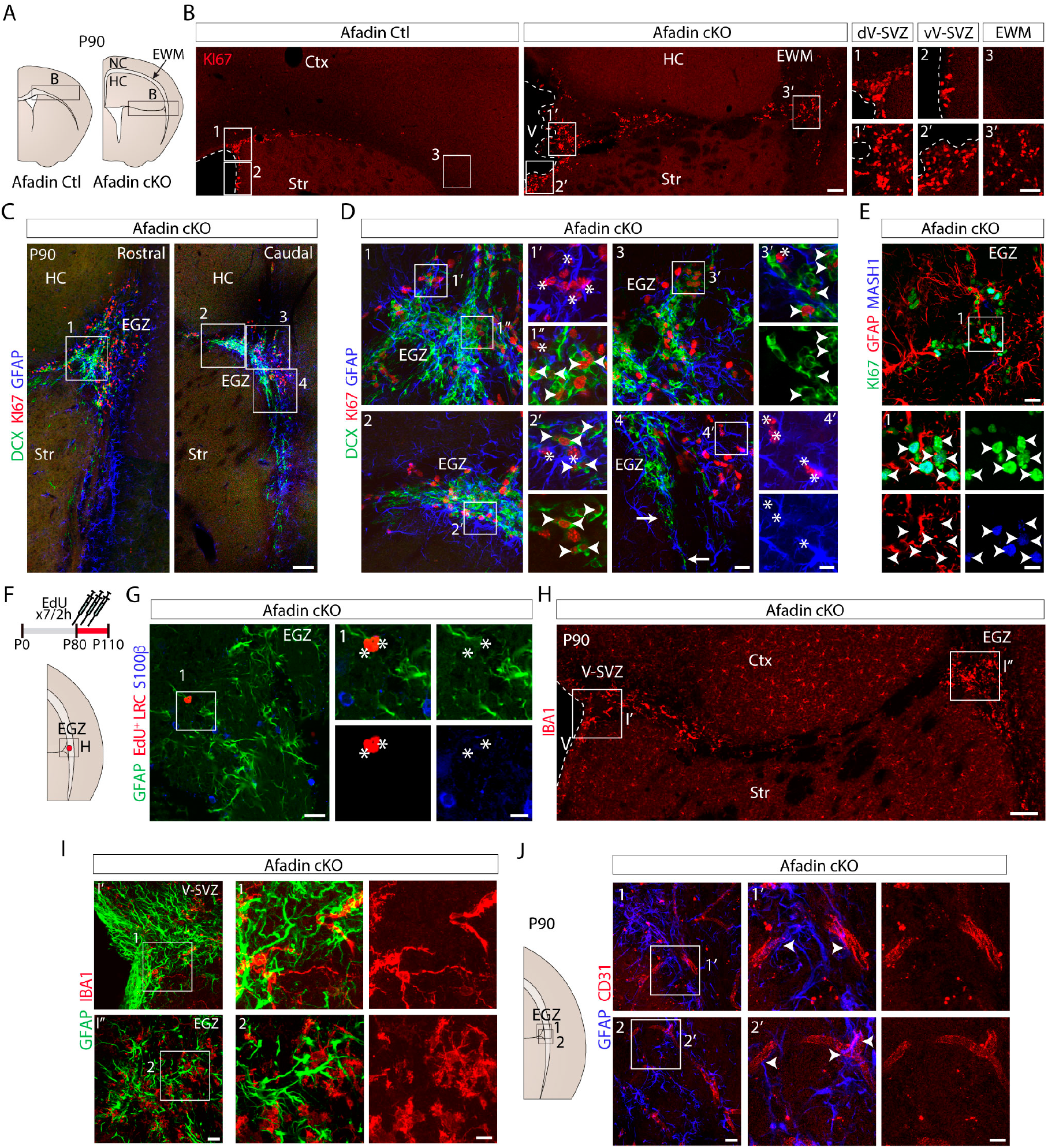
Developmental cortical remodeling gives rise to an ectopic neurogenic niche in the adult brain. **A**. Schematic of brain structures in *Afadin^Ctl^* and *Afadin^cKO^*mice. **B**. KI67 immunohistochemistry in coronal sections of control and mutant mice (P90), showing increased proliferation in the ventricular-subventricular zone (V-SVZ) (1’-2’) and cycling cells in ectopic regions of mutants (3’). **C**. Rostrocaudal panoramic views of the ectopic germinal zone (EGZ) in adult *Afadin^cKO^* mice immunostained for GFAP, KI67 and DCX. **D**. Higher magnification of boxed areas in C showing active neural stem cells (NSCs) (GFAP⁺Ki67⁺,*) and proliferative neuroblasts (DCX⁺Ki67⁺, arrowheads) in the EGZ. Arrows indicate possible migrating DCX^+^ cells. **E**. ASCL1⁺KI67⁺ transient amplifying progenitors (TAPs) in the EGZ. Arrowheads in the boxed area indicate double-labelled TAPs. **F**. Schematic of the EdU pulse–chase protocol used to identify label-retaining cells (LRCs). **G**. LRCs, including slowly dividing NSCs (EdU⁺GFAP⁺S100β⁻,*) are present in the EGZ of *Afadin^cKO^* mice. **H**. IBA1 immunohistochemistry reveals microglial cells in the EGZ with the typical ameboid morphology of the V-SVZ. **I**. The long processes of GFAP^+^ cells are in close apposition to IBA1^+^ microglial cells both in the V-SVZ (**I’**) and in the EGZ (**I’’**). **J.** CD31⁺ endothelial cells are closely associated with GFAP⁺ processes in the EGZ. Ctx, cortex; EWM, ectopic white matter; HC, heterocortex; NC, normocortex; Str, striatum; V, ventricle; d/vV-SVZ, dorsal/ventral ventricular– subventricular zone. Scale bars: **B**, **C**, **H** 100 µm; **B (1-3’)** 50 µm; **D (1-4)**, **E (1)**, **G**, **H (1-2)**, **I (1-2)** and **J (1-2)** 20 µm; **D (1’-4’**), **E (1’)**, **G (1)**, **I (1’-2’)** and **J (1’-2’)** 10 µm.

**Figure 2.**
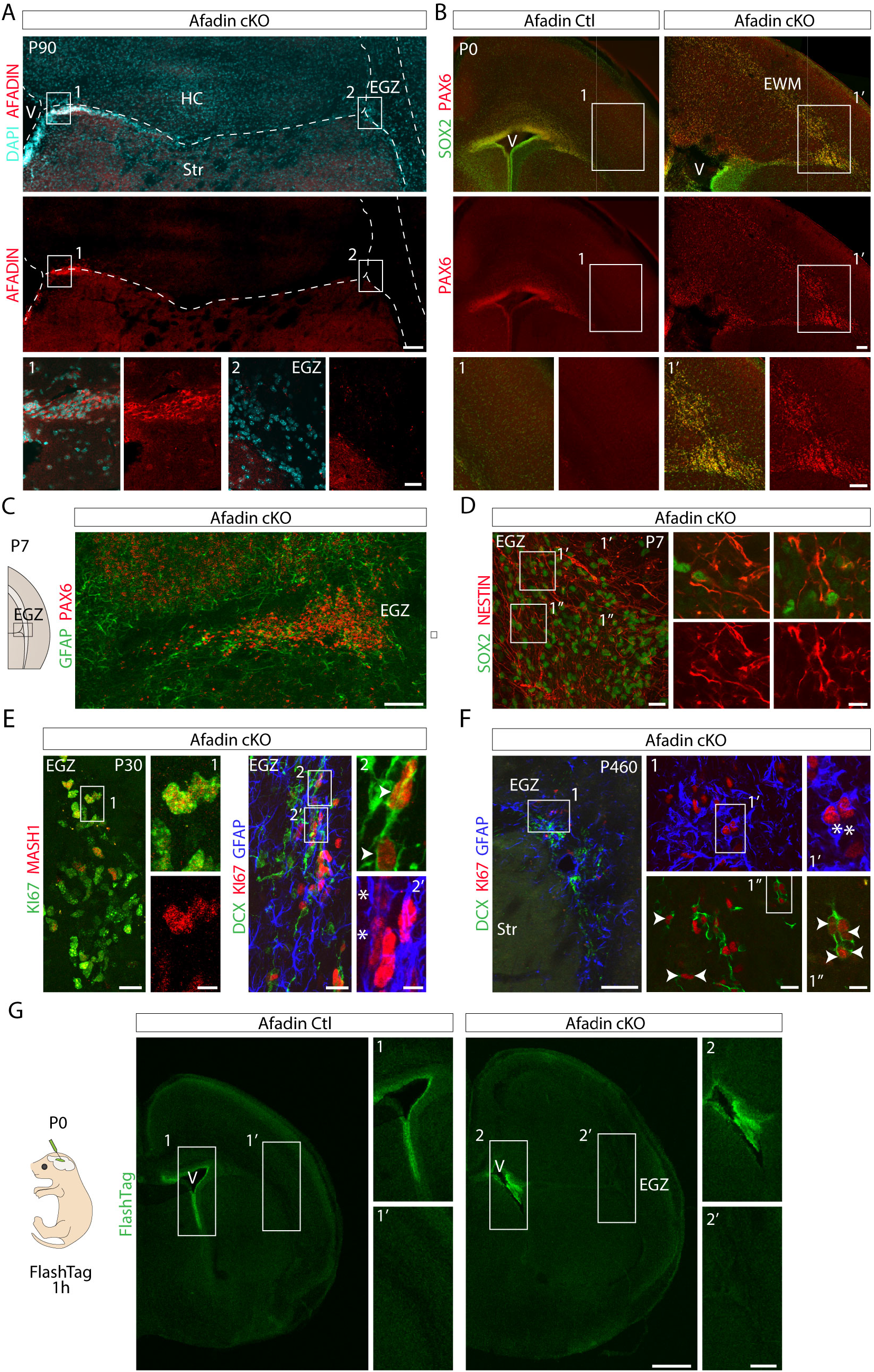
Ontogeny, long-term persistence, and ventricular independence of the ectopic niche. **A**. AFADIN expression is absent in EGZ cells, supporting their dorsal origin. **B**. PAX6 and SOX2 immunohistochemistry at P0 shows dispersion of radial glial cells (RGCs) throughout the mutant cortex, with prominent accumulation in the presumptive EGZ (arrowheads). **C**. PAX6 high and GFAP expression is detected in the EGZ at P7. **D**. Double immunohistochemistry of the two NSC markers SOX2 and NESTIN reveals their presence in the EGZ at P7. **E**. Proliferative TAPs (1’, ASCL1^+^KI67^+^), neuroblasts (2’, arrowheads, DCX^+^KI67^+^), and NSCs (2’’, *, GFAP^+^KI67^+^) are detected in the EGZ of young mice (P30). **F**. GFAP, KI67, and DCX labeling in aged *Afadin^cKO^* mice (P460) indicates persistent neurogenesis in EGZ. **G**. Intraventricular *FlashTag* labeling at P0.5 followed by 1 h survival of the pup labels only ventricular-adjacent cells in both control and mutant mice, indicating that the EGZ is disconnected from the ventricular surface in mutants. Ctx, cortex; EGZ, ectopic germinal zone; EWM, ectopic white matter; Str, striatum; V, ventricle. Scale bars: **A-C**, 100 µm; **D (1)**, **E (1, 2)**, **F (1)**, 20 µm; **D (1’-1’’)**, **E (1’, 2’, 2’’)**, **F (1’)**, 10 µm**; F (1’’)**, 5 µm; **G**, 500 µm; **G (1-2, 1’-2’)**; 200 µm.

**Figure 3.**
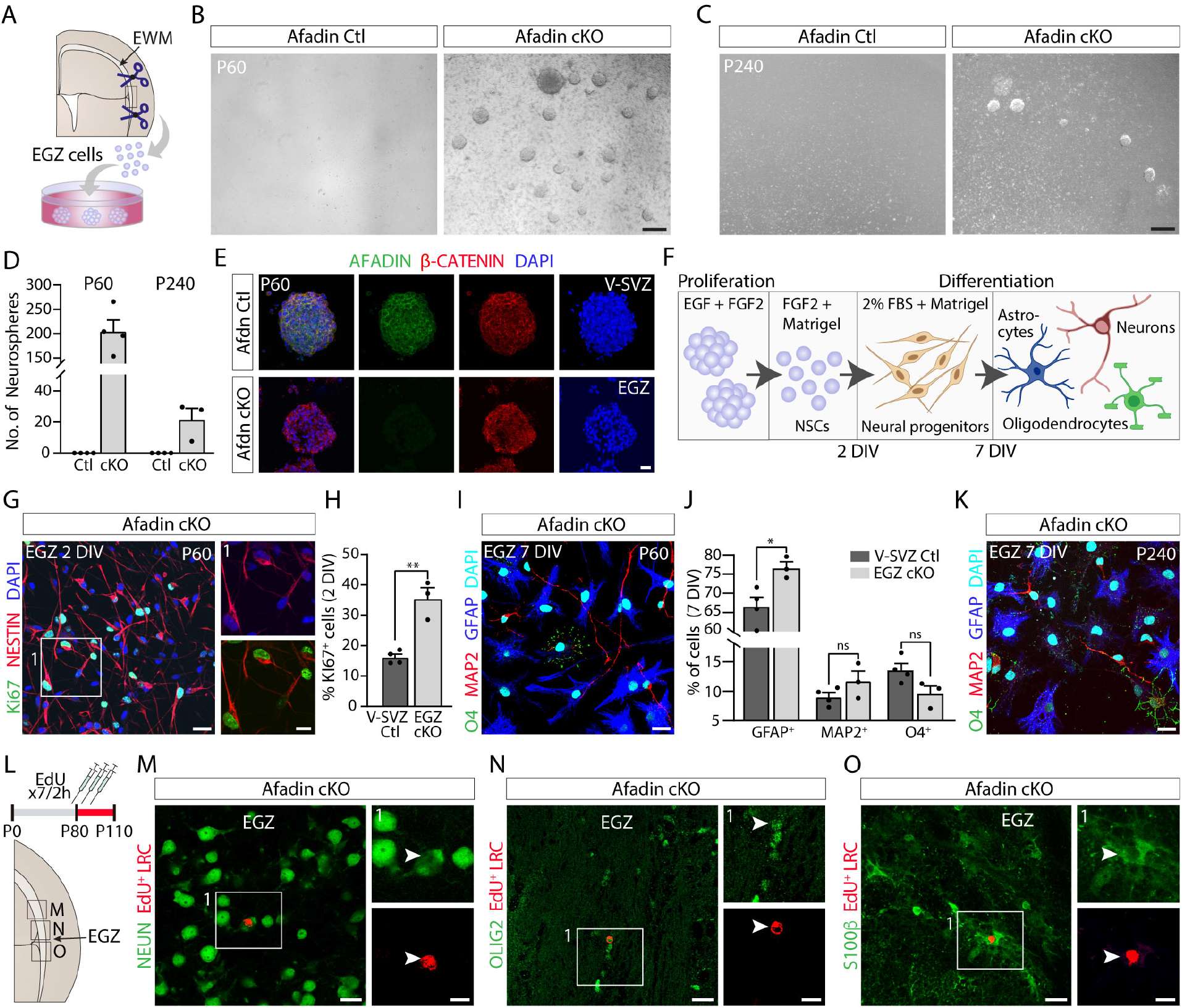
The ectopic neural stem cell niche harbors self-renewing, multipotent neural stem cells. **A**. Schematic of the experimental workflow for neurosphere assays from microdissected EGZ tissue of adult *Afadin^cKO^* brain and cortical tissue from controls, including tissue dissection, cell seeding, and *in vitro* proliferation and differentiation. **B-C.** Primary neurospheres formed exclusively from EGZ cultures of P60 (B) and P240 (C) mice. **D**. Quantification of B and C (mean ± s.e.m.; *n* = 4; statistical analysis was not performed as no neurospheres formed in control cultures). **E**. Control V-SVZ neurospheres express AFADIN and β-CATENIN, whereas EGZ-derived neurospheres lack AFADIN expression. **F**. Proliferative progenitors (NESTIN⁺KI67⁺) derived from EGZ neurospheres after 2 DIV under differentiation-promoting conditions. **G**. Quantification of NESTIN⁺KI67⁺ cells in cultures shown in F, relative to control V-SVZ neurospheres (mean ± s.e.m.; *n* = 3–4; \*\**p* = 0.0020; unpaired two-tailed Student’s *t*-test). **H**. Differentiated cells of EGZ-derived neurospheres after 7 DIV under differentiation-promoting conditions, immunostained for GFAP (astrocytes), MAP2 (neurons), and O4 (oligodendrocytes). **I**. Quantification of cell types shown in H, relative to control V-SVZ neurospheres (mean ± s.e.m.; *n* = 3–4; GFAP, \**p* = 0.0235; MAP2, *p* = 0.1694; O4, *p* = 0.0627; unpaired two-tailed Student’s *t*-test). **J**. GFAP, O4 and MAP2 immunocytochemistry in cells derived from EGZ neurospheres of aged *Afadin^cKO^* mice (P240) after 7 DIV. **K**. Schematic of the EdU pulse–chase protocol used to identify LRCs in the EGZ. **L-N.** EdU⁺ LRCs identified in the EGZ. Some EdU⁺ LRCs co-express markers of mature neurons (L, NEUN, arrowhead), oligodendrocytes (M, OLIG2, arrowhead) and astrocytes (N, S100β, arrowhead), demonstrating *in vivo* multipotency of EGZ-derived NSCs. DIV, days *in vitro*; EGZ, ectopic germinal zone; EWM, ectopic white matter; V-SVZ, ventricular–subventricular zone. Scale bars: **B**, **C**, 200 µm; **E**, **G**, **I**, **K**, **M–O**, 20 µm; **insets**, 10 µm.

We next asked whether the EGZ represents a neurogenic niche rather than a simple ectopic proliferative domain by characterizing its cellular organization. We identified actively dividing NSCs^28^ (GFAP⁺KI67⁺; Fig. 1D, asterisks) closely associated with DCX^+^ immature neurons^29^, a subset of which remained proliferative (DCX⁺KI67⁺; Fig. 1D, arrowheads). Notably, several DCX^+^ neurons directed their leading processes towards the external capsule, suggesting migratory behavior (Fig. 1C-D, arrows). Co-immunostaining for ASCL1⁺ and KI67⁺ further identified proliferating transit-amplifying progenitors (TAPs)^30^ (Fig. 1E, arrowheads). qPCR analysis further demonstrated increased expression of the neurogenic markers *Pax6*, *Ascl1*, and *Dcx* in mutant EGZ tissue relative to wild-type cortex (Extended Data Fig. 1C). Furthermore, long-term EdU labeling revealed EdU⁺ label-retaining cells (LRCs; Fig. 1F) in this germinal zone, including slowly dividing NSCs (EdU⁺GFAP⁺S100β⁻ cells; Fig. 1G).

We then assessed whether the EGZ also contains additional cellular and structural components characteristic of an NSC niche. The EGZ contained IBA1⁺ microglia with an amoeboid morphology reminiscent of V-SVZ microglia^31^ and distinct from the ramified microglia observed in the neocortex (Fig. 1H). IBA1/GFAP co-immunostaining revealed that these amoeboid microglia were closely apposed to the elongated processes of GFAP⁺ cells resembling the pattern found in the V-SVZ (Fig. 1I). In addition, a CD31⁺ vascular network was closely associated with GFAP⁺ processes within the EGZ, resembling the vascular organization described in the V-SVZ ^9,14^ (Fig. 1J). Thus, beyond housing the major neurogenic cell populations, the EGZ displays key cellular and structural features of an organized NSC niche.

These observations reveal that developmental disruption of cortical architecture can durably reconfigure the adult NSC landscape *in vivo*, generating an ectopic domain that comprises cellular and structural components of a neurogenic niche

### Perinatal pallial progenitors seed an ectopic niche that endures into aging beyond the ventricular surface

Having established the presence of an ectopic neurogenic niche in adult brains, we next investigated whether this domain emerges during embryonic and early postnatal development. We first examined AFADIN expression within the EGZ. Cells in this region lacked AFADIN protein (Fig. 2A), consistent with their derivation from *Afadin*-deficient dorsal pallial territories. We therefore asked whether the EGZ could originate from the delamination of pallial radial glial cells (RGCs) previously observed during embryonic cortical development following *Afadin* loss^22^.

To test this possibility, we examined the early postnatal distribution of pallial progenitors using PAX6 immunohistochemistry^32^. At P0, PAX6⁺ cells accumulated beneath the normocortex, along the trajectory of projection neuron axons that form the EWM (Fig. 2B). This accumulation was particularly prominent in the lateralmost EWM, at the position subsequently occupied by the EGZ. These PAX6⁺ cells co-expressed SOX2 (Fig. 2B), supporting their identity as undifferentiated pallial progenitors. Together with the absence of AFADIN in the adult EGZ, these findings indicate that delaminated pallial progenitors seed the ectopic domain during the perinatal period.

We next examined the early postnatal organization of this ectopic progenitor population. At P7, PAX6⁺ cells remained concentrated within the EGZ, which had become a GFAP-rich domain (Fig. 2C). The EGZ also contained NESTIN^+^SOX2^+^ cells, both markers of NSCs since early ages^33,34^, further supporting the establishment of an NSC-like population at this ectopic domain (Fig. 2D). By P30, the EGZ contained the full complement of neurogenic cell populations observed in the adult niche, including NSCs, TAPs, and neuroblasts (Fig. 2E). Interestingly, this organization remained still evident in aged animals at P460 (Fig. 2F), demonstrating its long-term persistence.

Given the ectopic location of the EGZ within the lateralmost EWM, away from the ventricular surface, we next asked whether tissue disorganization nevertheless provides this domain with direct access to CSF-derived signals. To address this possibility, we performed intraventricular *FlashTag* labeling at P0, when PAX6⁺ progenitors begin to accumulate within the lateralmost EWM, marking the emergence of the future EGZ, and the surrounding parenchyma remains relatively loosely organized. *FlashTag* labels cells directly contacting the ventricular surface^35^ and can reveal dissemination of ventricular contents into the adjacent parenchyma. One hour after injection, *FlashTag* labeling was restricted to cells lining the ventricular surface in both control and mutant brains, with no signal detected in the surrounding parenchyma, including the EGZ (Fig. 2G). Thus, the EGZ is not directly exposed to the ventricular lumen during its formation, arguing that this ectopic niche is established independently of direct CSF contact.

Overall, these findings support a model in which developmental delamination of pallial progenitors is an initiating event through which tissue disorganization gives rise to a spatially ectopic neurogenic niche that endures into aging without direct ventricular contact.

### Ectopic NSCs retain self-renewal and multipotency with age

Having established the cellular organization of the ectopic niche and its origin, we next asked whether it supports functional NSCs with the defining properties of self-renewal and multipotency. To address this, we performed neurosphere assays using microdissected EGZ tissue from young adult (P60) and middle-aged (P240) *Afadin^cKO^* mice (Fig. 3A) following published protocols^36,37^. Cortical tissue from control mice was used as a negative control. After 7 days *in vitro* (DIV) in mitogen-enriched medium containing epidermal growth factor (EGF) and fibroblast growth factor 2 (FGF2), control samples failed to generate neurospheres, whereas EGZ-derived cultures robustly formed primary neurospheres at both ages (Fig. 3B-C). Quantification revealed that primary neurosphere numbers in young mutants were comparable to those derived from the control V-SVZ tissue^38,39^, and declined with age (Fig. 3D), mirroring the age-dependent decrease observed in V-SVZ-derived cultures^40^. Secondary neurospheres derived from the EGZ lacked AFADIN immunoreactivity, confirming their dorsal origin (Fig. 3E).

We next examined the differentiation potential of EGZ-derived neural stem cells using established *in vitro* differentiation assays^36,37^. Neurospheres were dissociated and plated under adherent conditions in FGF2-containing medium to promote neural progenitor identity (Fig. 3A). At 2 DIV, most cells were NESTIN^+^ and exhibited neural progenitor morphology (Fig. 3F). KI67 immunostaining showed a significantly higher proliferation rate in EGZ-derived progenitors compared with control V-SVZ cultures (Fig. 3F–G), indicating that they retain robust proliferative capacity after isolation.

To determine whether EGZ-derived stem cells were multipotent, cultures were subsequently switched to serum-containing to induce terminal differentiation (Fig. 3A). After 5 additional days, immunostaining for lineage markers (MAP2, neurons; GFAP, astrocytes; O4, oligodendrocytes)^41–43^ demonstrated the generation of the three neural lineages from EGZ-derived NSCs of young adults (P60) (Fig. 3H). Quantification revealed a slight increase in the astrocyte fraction in EGZ-derived cultures, whereas neuronal and oligodendrocyte fractions were comparable to those in wild-type V-SVZ-derived NSCs (Fig. 3I). Similarly, neurospheres derived from P240 EGZ tissue also generated neurons, astrocytes, and oligodendrocytes (Fig. 3J), demonstrating preservation of multipotency at advanced ages.

To assess whether EGZ NSCs also generate multiple neural lineages *in vivo*, we analyzed the long-term EdU labeling assays described above (Fig. 3K) to look for EdU^+^ cells in the vicinity of the ectopic niche. Although EdU incorporation alone cannot definitively establish cellular origin, several observations strongly support an EGZ origin, including the adult-onset labeling, the distance from the V-SVZ, and the compact organization of the surrounding tissue. EdU⁺ cells co-expressing neuronal (NEUN⁺)^44^, astrocytic (S100β)^45^, or oligodendrocyte (OLIG2)^46^ markers were identified near the EGZ (Fig. 3L-N), indicating that EGZ NSCs generate all three major neural lineages *in vivo*.

Together, these findings demonstrate that the ectopic niche harbors functional NSCs that retain the defining properties of stem cells both *in vitro* and *in vivo*.

### Developmental perturbation reshapes endogenous neural stem cell niche architecture

The formation of a persistent ectopic neural stem cell niche raised the question of how the native V-SVZ niche is affected by the same developmental perturbation. Because EGZ NSCs originate from dorsal progenitors that normally contribute to the canonical adult V-SVZ and given the established role of *Afadin* in regulating apico-basal polarity^47^, we asked how *Afadin* deletion alters the adult V-SVZ architecture.

Whole-brain MRI confirmed previously described structural abnormalities in *Afadin^cKO^*^22^, including double-cortex formation, cortical enlargement, and commissural defects such as agenesis of the corpus callosum (Fig. 4A). MRI also revealed marked rostrocaudal changes in ventricular size, volume, and shape (Extended Data Fig. 2A–B), indicating remodeling of the ventricular architecture that supports the endogenous V-SVZ niche.

**Figure 4.**
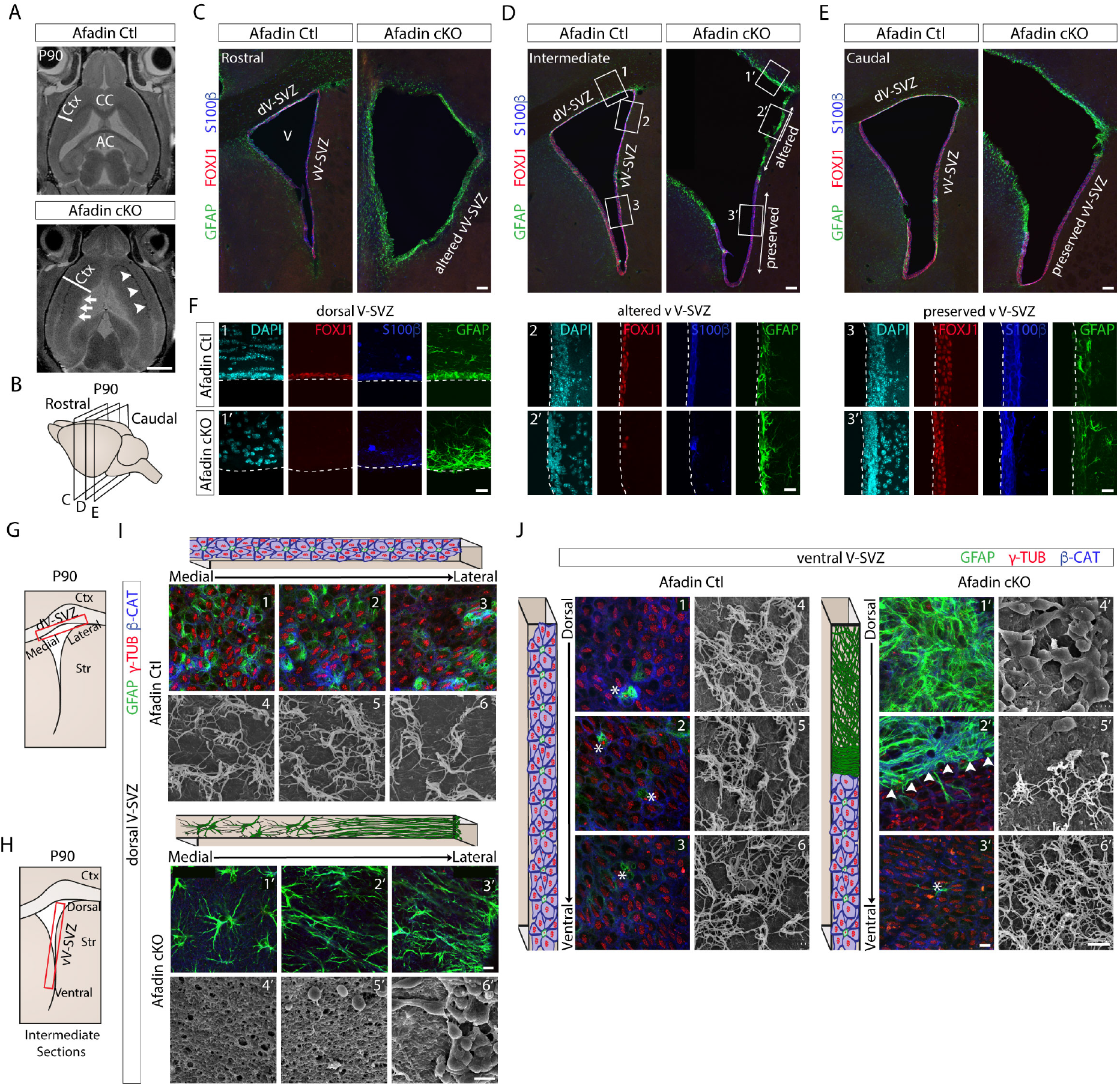
The endogenous V-SVZ niche undergoes extensive structural reorganization. **A**. Magnetic resonance imaging (MRI) of adult control and *Afadin^cKO^* brains reveals an irregular ventricular surface in mutants (arrows), together with cortical enlargement and double-cortex formation (arrowheads). **B**. Schematic showing the coronal levels analyzed along the rostrocaudal axis of the adult brain ventricular surface. **C-E.** Immunohistochemistry for GFAP, FOXJ1, and S100β reveals loss of ependymal markers and expansion of GFAP⁺ cells in the dorsal V-SVZ of the mutants. The mutant ventral V-SVZ shows differential alterations across the rostrocaudal axis, being more prominent rostrally (C) than caudally (E). **F**. Higher magnification of the boxed areas in D showing ependymal cells loss in the dorsal V-SVZ (1-1’) and the altered ventral V-SVZ (2-2’), and preservation of ependymal cell markers in unaffected ventral region (3-3’). **G-H**. Schematic of dorsal and ventral V-SVZ *en face* dissections for immunohistochemical and scanning electron microscopy (SEM) analyses. **I**. 1-3 and 1’-3’ show loss of γ-TUBULIN⁺ multiciliated cells and replacement by GFAP⁺ cells in *Afadin^cKO^*mice. 4-6 and 4’-6’ confirm, by SEM, the cilia loss and exposure of cell bodies in the dorsolateral region in *Afadin^cKO^* mice. **J**. 1-3 and 1’-3’ identify an altered ventral region lacking multiciliated cells and a preserved ventral region retaining ependymal cells and pinwheel organization (*), separated by a dense GFAP⁺ band (arrowheads). 4-6 and 4’-6’ confirm, by SEM, immunohistochemical analysis and reveal increased exposure of cell bodies in the altered ventral V-SVZ. AC, anterior commissure; CC, corpus callosum; Ctx, cortex; d/vV-SVZ, dorsal/ventral ventricular–subventricular zone. Scale bars: **A**, 5 mm; **C-E**, 100 µm; **F**, 20 µm; **I** and **J (1-3**, **1’-3’)**, 10 µm; **I** and **J (4-6**, **4’-6’)**, 5 µm.

Immunohistochemical analysis across rostrocaudal levels revealed profound disruption of the dorsal ventricular wall in *Afadin^cKO^* mice, characterized by reduced cell density, loss of ependymal markers FOXJ1 and S100β^13,48,49^, and increased expression of the astrocytic and NSC marker GFAP (Fig. 4B-F). However, abnormalities also extended into adjacent ventral V-SVZ territories where *Afadin* expression is preserved, indicating that developmental disruption occurs beyond the primary domain of *Afadin* deletion. These changes displayed a rostrocaudal gradient: rostral regions showed severe ventricular wall disruption, whereas caudal territories retained relatively preserved architecture (Fig. 4C-E). Intermediate regions contained both altered and preserved domains, with loss of FOXJ1 and S100β expression restricted to the upper ventral V-SVZ while adjacent lower regions maintained normal ependymal organization (Fig. 4D; Fig. 4F). The spatial association between cortical enlargement and ventral V-SVZ disruption was further supported by MRI measurements, as the greatest cortical expansion occurred at rostral levels where ventricular remodeling was most pronounced (Extended Data Fig. 2C-E).

Immunohistochemistry and scanning electron microscopy (SEM) of *en face* preparations further confirmed loss of normal dorsal ventricular organization. In control mice, β-CATENIN and γ-TUBULIN staining showed the characteristic honeycomb organization of multiciliated ependymal cells^13^ (Fig. 4I, 1-3), which was corroborated by SEM (Fig. 4I, 4-6). In contrast, mutant mice lacked this organization, with GFAP⁺ cells displaying abnormal morphologies (Fig. 4I, 1’-3’) and SEM revealing a smoother ventricular surface with exposed underlying cell bodies (Fig. 4I, 4’-6’). Similar alterations were observed in affected ventral V-SVZ regions, where abnormal GFAP⁺ cells replaced the typical pinwheel organization of the adult niche (Fig. 4J, 1-3). Notably, a dense GFAP⁺ boundary demarcated altered and preserved ventral territories (Fig. 4J, 2’, arrowheads). Consistently, SEM analysis revealed smooth ventricular surfaces with prominent exposure of underlying cell bodies specifically within altered ventral domains (Fig. 4J, 4’-5’), whereas densely ciliated surfaces were maintained in control and neighboring preserved ventral regions (Fig. 4J, 4-6, 6’).

**Figure 5.**
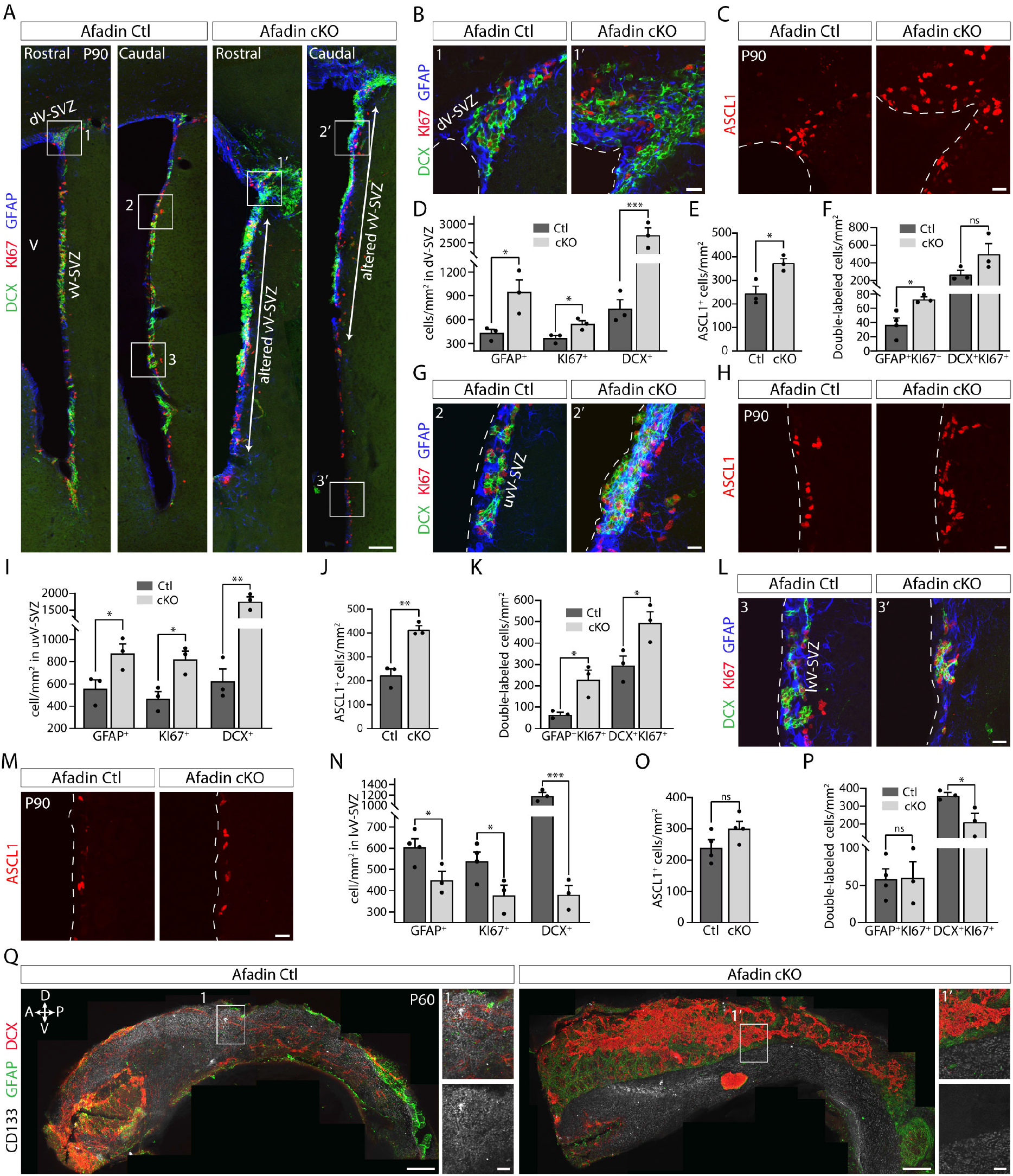
V-SVZ niche reorganization produces distinct neurogenic states. **A.** Immunostaining for GFAP, KI67 and DCX in the V-SVZ of *Afadin^Ctl^* and *Afadin^cKO^* mice (P90) along the rostrocaudal axis. Altered regions in mutant mice mirror the structural alterations of V-SVZ. **B.** Higher magnification of boxed areas 1-1’ in A displaying the dorsal V-SVZ in control and mutant mice. **C.** ASCL1 immunohistochemistry in the dorsal V-SVZ of both genotypes. **D–F.** Quantification of the labeled cell populations shown in B-C (mean ± s.e.m.; n = 3–4; GFAP⁺, \**p* = 0.0339; KI67⁺, \**p* = 0.0307; DCX⁺, \*\*\**p* = 0.0010; ASCL1⁺, \**p* = 0.0245; GFAP⁺KI67⁺, \**p* = 0.0234; DCX⁺KI67⁺, *p* = 0.1421; unpaired two-tailed Student’s *t*-test). **G.** Higher magnification of boxed areas 2-2’ in A displaying the altered ventral V-SVZ region in mutants and an equivalent location in control mice. **H.** ASCL1⁺ progenitors in the ventral V-SVZ at the locations shown in G. **I–K.** Quantification of the labeled cell populations shown in G-H (mean ± s.e.m.; n = 3; GFAP⁺, \**p* = 0.0482; KI67⁺, \**p* = 0.0198; DCX⁺, \*\**p* = 0.0030; ASCL1⁺, \*\**p* = 0.0036; GFAP⁺KI67⁺, \**p* = 0.0210; DCX⁺KI67⁺, \**p* = 0.0394; unpaired two-tailed Student’s *t*-test). **L.** Higher magnification of boxed areas 3-3’ in A showing a preserved ventral V-SVZ in mutants and control mice. **M.** ASCL1⁺ progenitors in ventral V-SVZ locations shown in L. **N–P.** Quantification of the labeled cell populations shown in L-M (mean ± s.e.m.; n = 3–4; GFAP⁺, \**p* = 0.0371; KI67⁺, \**p* = 0.0489; DCX⁺, \*\*\**p* = 0.0004; ASCL1⁺, *p* = 0.1137; GFAP⁺KI67⁺, *p* = 0.9534; DCX⁺KI67⁺, \**p* = 0.0390; unpaired two-tailed Student’s *t*-test). **Q**. Whole-mount V-SVZ preparations (P60) showing accumulation of DCX⁺ cells at the ventricular surface, above GFAP⁺ cells in regions lacking CD133⁺ cilia in *Afadin^cKO^* mice. A, anterior; D, dorsal; P, posterior; V, ventricle; d/vV-SVZ, dorsal/ ventral ventricular–subventricular zone. Scale bars: **A**, **Q (1-1’)**, 100 µm; **B**, **C**, **G**, **H**, **L**, **M**, 20 µm; **Q**, 300 µm.

**Figure 6.**
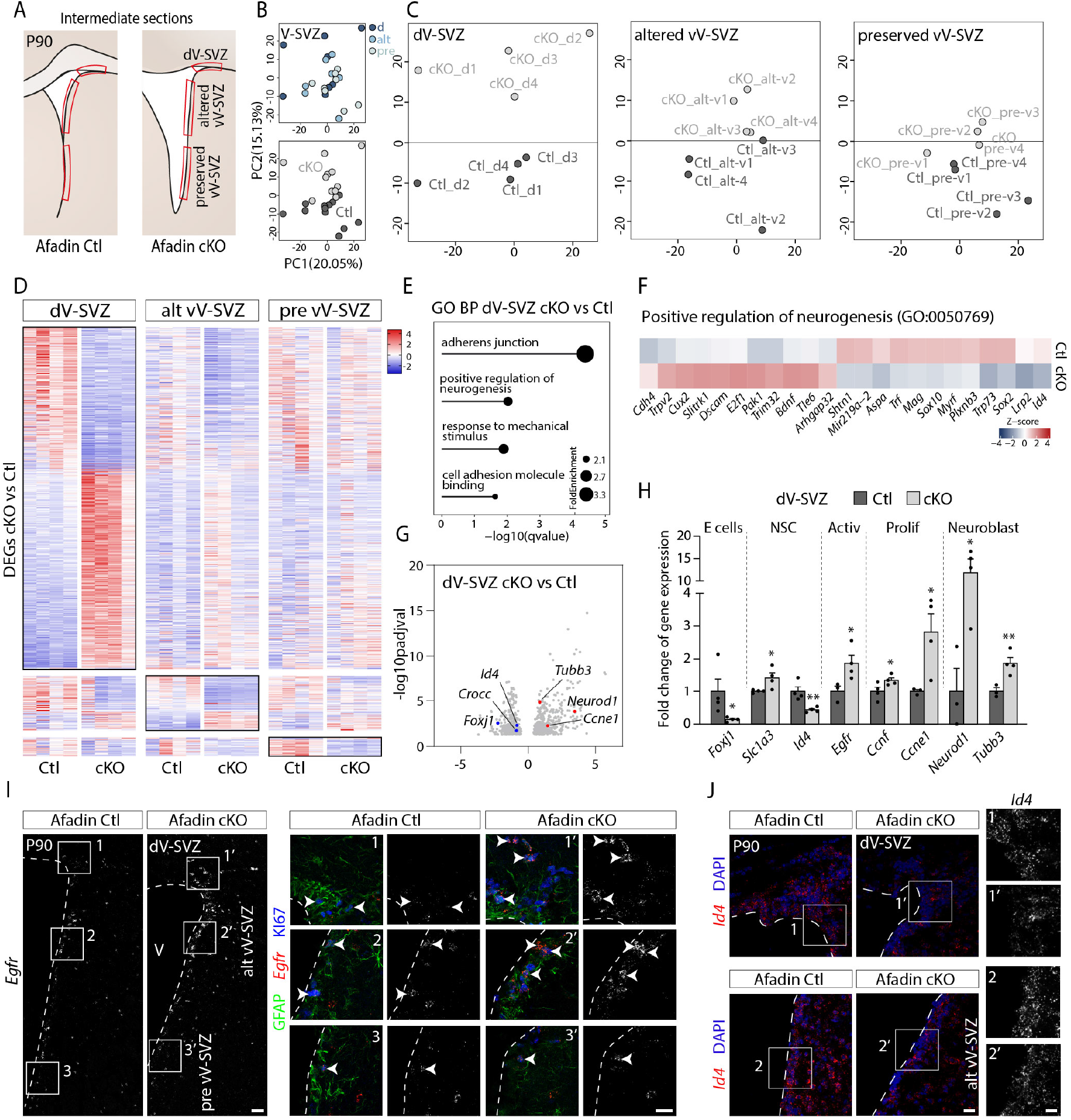
Transcriptional profiling reveals distinct molecular states across the adult V-SVZ niche. **A**. Schematic of intermediate coronal sections from adult control and *Afadin^cKO^* mice displaying the microdissected dorsal (d), altered ventral (alt-v) and preserved ventral (pre-v) V-SVZ subregions used for bulk RNA-seq. **B**. Principal component analysis (PCA) of bulk RNA-seq data (n = 4 per genotype) shows segregation primarily by genotype rather than anatomical region. Principal components 1 and 2 are shown. **C**. Region-specific PCA plots reveal strongest genotype-dependent separation in the dorsal V-SVZ, followed by the altered and preserved ventral domains. **D.** Heatmap of Z-score-scaled expression of differentially expressed genes (DEGs) identified between control and mutant mice across V-SVZ subregions. **E**. Gene Ontology (GO) analysis detects enrichment of biological processes in the dorsal V-SVZ of mutant mice, including adherens junction organization, cell adhesion, response to mechanical stimulus, and positive regulation of neurogenesis. Dot size indicates fold enrichment. **F**. Heatmap of DEGs associated with the GO term *Positive regulation of neurogenesis* (GO:0050769). **G**. Volcano plot of DEGs in the dorsal V-SVZ of *Afadin^Ctl^* vs *Afadin^cKO^*mice. The x-axis represents the average fold change between *Afadin^cKO^* and control mice, and the y-axis shows the statistical significance (-log10FDR). Representative downregulated (blue; *Id4*, *Foxj1*, *Crocc*) and upregulated (red; *Ccne1*, *Neurod1*, *Tubb3*) genes are indicated. **H**. qPCR validation of selected DEGs in dorsal V-SVZ tissue (mean ± s.e.m.; n = 3-4; *Foxj1*, \**p* = 0.0352; *Slc1a3*, \**p* = 0.0194; *Id4*, \*\**p* = 0.0052; *Egfr*, \**p* = 0.0453; *Ccnf*, \**p* = 0.03176; *Ccne1*, \**p* = 0.0205; *Neurod1*, \**p* = 0. 0317; *Tubb3*, \*\**p* = 0.0061; unpaired one-tailed Student’s *t*-test). **I-J**. RNAscope combined with KI67 and GFAP immunostaining confirms increased *Egfr* expression (arrowheads) in proliferative cells (I) and reduced *Id4* expression (J) in the dorsal and altered ventral V-SVZ in *Afadin^cKO^*mice. d/alt/pre vV-SVZ, dorsal/altered ventral/preserved ventral ventricular-subventricular zone; V, ventricle. Scale bars: **I**, 100 µm; **I (1-3’)**, **J**, 20 µm; **J (1-2’)**, 10 µm.

The regional differences observed in adult V-SVZ remodeling suggested that distinct developmental mechanisms may underlie the loss of mature ependymal organization. We therefore analyzed ependymal specification and differentiation during the early postnatal period, when the ventricular lining is established^50–52^ (P0–P7; Extended Data Fig. 3A-D). In control mice, FOXJ1⁺CD133⁺ ependymal cells progressively appeared throughout the ventricular walls. In *Emx1*-derived dorsal territories of *Afadin^cKO^* mice, FOXJ1 expression was absent at all analyzed stages, indicating failure to acquire ependymal identity following *Afadin* deletion (Extended Data Fig. 3B-D). Conversely, affected ventral territories, where *Afadin* expression is preserved, initiated ependymal specification normally, as indicated by comparable FOXJ1 expression at P0 (Fig. 3B), but failed to complete terminal differentiation, resulting in loss of mature FOXJ1⁺CD133⁺ ependymal cells at later stages (Extended Data Fig. 3 C-D).

The absence of mature ependymal cells was not explained by increased apoptosis, as activated CASP3 immunoreactivity remained scarce near the ventricular wall (Extended Data Fig. 3E-G). Instead, affected regions accumulated GFAP⁺ cells with features associated with immature radial glia-like states^48^. In the adult dorsal V-SVZ, including the neurogenic wedge region, a specialized domain along the dorsal ventricular wall that contributes to adult neurogenesis^53^, as well as in altered ventral V-SVZ territories lacking mature CD133⁺S100β⁺ ependymal cells, GFAP⁺ cells expressing high levels of BLBP were detected in *Afadin^cKO^* mice, whereas BLBP expression was low in the corresponding regions of control mice (Extended Data Fig. 3H). Consistent with persistence of an immature progenitor-like state, both remodeled regions also exhibited increased NESTIN expression (Extended Data Fig. 3I), a marker also associated with activated adult NSCs ^54–56^.

Thus, developmental *Afadin* loss reveals how tissue disorganization reshapes V-SVZ architecture across both directly affected dorsal and adjacent ventral territories. This rostrocaudally graded remodeling parallels cortical expansion and features failed ependymal specification dorsally, impaired maturation ventrally, and retention of radial glia-like cells.

### **V-** SVZ niche reorganization alters adult neurogenic activity and neuronal integration

Having established that developmental perturbation simultaneously generates an ectopic niche and remodels the endogenous V-SVZ, we next asked how these structural changes influence neurogenic activity and the integration of newborn neurons in the adult brain.

Immunohistochemical analysis of distinct neurogenic populations throughout the V-SVZ revealed a coordinated expansion of neurogenic populations in the dorsal zone and differential responses across ventral domains of *Afadin^cKO^* mice (Fig. 5A), mirroring the structural defects previously described along the rostrocaudal axis (Fig. 4C–E). Within the dorsal neurogenic wedge region, *Afadin^cKO^* mice exhibited increases in GFAP^+^ cells, KI67^+^ proliferating cells, and DCX^+^ neuroblasts compared with controls (Fig. 5B; Extended Data Fig. 4A). ASCL1 immunostaining further revealed expansion of TAPs (Fig. 5C), and quantification confirmed significant increases in these populations (Fig. 5D–E). The number of GFAP^+^KI67^+^ cells was also increased, consistent with enhanced NSC activation. Proliferative neuroblasts (DCX^+^KI67^+^) tended to be more numerous in mutants, although this difference did not reach statistical significance (Fig. 5F).

A similar expansion of neurogenic populations was observed in structurally altered regions of the ventral V-SVZ. These territories contained increased numbers of GFAP^+^, KI67^+^, DCX^+^, and ASCL1^+^ cells relative to controls (Fig. 5G–J; Extended Data Fig. 4B), together with elevated numbers of GFAP^+^KI67^+^ NSCs and DCX^+^KI67^+^ neuroblasts (Fig. 5K). In contrast, structurally preserved ventral regions displayed the opposite phenotype. Numbers of GFAP^+^, KI67^+^, and DCX^+^ cells were reduced, whereas ASCL1^+^ cell numbers were unchanged (Fig. 5L–P; Extended Data Fig. 4C). GFAP^+^KI67^+^ NSCs were not significantly different, whereas DCX^+^KI67^+^ neuroblasts were significantly reduced (Fig. 5P).

To determine how these changes were spatially organized across the ventricular surface, we analyzed V-SVZ whole-mounts immunostained for multiple cellular markers (Fig. 5Q). These analyses revealed widespread accumulation of DCX^+^ cells along the apical surface of the mutant V-SVZ, largely within CD133^−^ areas devoid of ependymal cells (Fig. 4Q, 1′). Three-dimensional reconstruction confirmed that DCX^+^ cells formed a superficial layer above GFAP^+^ cells in the altered ventral V-SVZ (Extended Data Fig. 4D; Movie), identifying the nature of the exposed cell bodies observed by SEM (Fig. 4H–I). Notably, a substantial fraction of these apically located DCX^+^ cells co-expressed KI67, indicating that many remained proliferative at the adult ages analyzed (Extended Data Fig. 4E, arrows).

We next asked whether this altered neurogenic output was accompanied by defects in neuronal migration and integration. Given the established roles of *Afadin* in neuronal migration^22,57^, we examined the rostral migratory stream (RMS) in adult *Afadin^Ctl^* and *Afadin^cKO^* mice using DCX to label migrating neurons (Extended Data Fig. 5A). Whereas control mice displayed a compact RMS with tightly aligned DCX^+^ cells, *Afadin^cKO^* mice exhibited an expanded and disorganized RMS, particularly within the intermediate segment, where DCX^+^ cells deviated along aberrant upper and lower trajectories and appeared dispersed into neighboring tissues (Extended Data Fig. 5A).

Reconstruction of binarized RMS from serial DCX-immunostained sections confirmed a significant increase in RMS area accompanied by reduced DCX^+^ pixel density (Extended Data Fig. 5B–D), consistent with impaired organization of migrating neuronal chains. Despite this reduced density, total DCX^+^ signal was increased in mutants, indicating that enhanced neuronal production was uncoupled from efficient migratory organization (Extended Data Fig. 5E). Orientation analysis further supported this defect: whereas DCX^+^ elements in controls were predominantly aligned around 0°, reflecting directed migration toward the olfactory bulb (OB), mutant RMS lacked a dominant orientation and showed a broad distribution of angles (Extended Data Fig. 5F–H).

We then examined whether these migratory abnormalities were associated with altered neuronal accumulation within the OB. Although overall laminar organization was preserved, the OB was significantly smaller and exhibited an expanded RMS entry zone with reduced cell density in *Afadin^cKO^* mice (Extended Data Fig. 5I–K). Consistent with impaired organization of the migratory stream, DCX^+^ cells were more broadly distributed within the OB-RMS region of mutant mice and showed reduced signal intensity (Extended Data Fig. 5L–N). Finally, to directly assess long-term neuronal incorporation into the OB, we performed the EdU LRC assay described previously (Fig. 1F). Quantification of coronal OB sections revealed a significant reduction in EdU^+^ LRCs in *Afadin^cKO^* mice compared with controls, both globally and within the granular cell layer (GCL) (Extended Data Fig. 5O–P).

Collectively, these findings demonstrate that developmental remodeling of the V-SVZ has lasting functional consequences for adult neurogenesis. Although neurogenic activity is enhanced in structurally remodeled domains, this increased output is uncoupled from efficient neuronal migration and long-term incorporation into the OB.

### Developmental remodeling establishes distinct transcriptional states within the adult V-SVZ niche

Because adhesion-associated proteins not only provide structural organization but also function as signaling platforms that regulate intracellular pathways and gene expression programs^58,59^, we asked whether early dorsal *Afadin* loss induces transcriptional alterations in the adult V-SVZ. Given that structural and proliferative remodeling extended beyond the primary dorsal domain of *Afadin* loss, we microdissected dorsal, altered ventral, and preserved ventral V-SVZ subregions from *Afadin^cKO^* mice and their respective controls (Fig. 6A). Intermediate rostrocaudal section in which altered (upper-ventral) and preserved (lower-ventral) domains could be visualized in the same section were used for microdissection (Fig. 4D). The accuracy of V-SVZ microdissection was confirmed by qPCR enrichment of neurogenic markers, including *Nestin* and *Ascl1*, relative to whole brain tissue (Extended Data Fig. 6A).

Bulk RNA-sequencing (RNA-seq) was performed on microdissected samples from the three V-SVZ regions. Principal component analysis (PCA) identified genotype as the major source of variation, while preserving regional distinctions among V-SVZ domains (Fig. 6B). When each region was analyzed independently, the strongest genotype-dependent separation was observed in the dorsal V-SVZ, followed by the altered and preserved ventral regions (Fig. 6C). Differential-expression analysis revealed a graded transcriptional response, with 1151, 142 and 56 differentially expressed genes in the dorsal, altered ventral, and preserved ventral domains, respectively (Fig. 6D).

Gene Ontology (GO) analysis of biological processes identified statistically significant differences between mutants and controls primarily in the dorsal V-SVZ of *Afadin^cKO^* mice, consistent with its stronger transcriptional shift. Enriched GO terms included categories related to loss of adhesion, such as adherens junction organization, cell adhesion molecule binding, as well as response to mechanical stimulus (Fig. 6E; Extended Fig. 6B). GO analysis also revealed enrichment of positive regulation of neurogenesis (Fig. 6E-F), in agreement with our histological findings. In line with this, dorsal V-SVZ samples from *Afadin^cKO^* mice showed downregulation of quiescence-associated regulators (e.g., *Id4*^60^) and ependymal-associated genes (e.g., *Foxj1*^48^*; Crocc* ^61^) (Fig. 6G). Conversely, transcripts associated with cell-cycle progression (e.g., *Ccne1*^62^) and early neuronal differentiation (e.g., *Neurod1*, *Tubb3* ^63,64^) were upregulated (Fig. 6G). These and additional transcriptional changes were independently validated by qPCR (Fig. 6H).

Despite the stronger transcriptional response in the dorsal V-SVZ, the altered ventral domain displayed similar cellular phenotypes (Fig. 5), raising the possibility that anatomically distinct regions converge on shared molecular states. To determine whether these regions shared subtle or cell-type–specific transcriptional signatures, we analyzed a curated gene set enriched in distinct V-SVZ subpopulations, including quiescent cells/astrocytes, NSCs, activated NSCs, TAPs, and neuroblasts^55,65^ across the three regions. The altered ventral V-SVZ transcriptomic profile more closely resembled that of the dorsal region than that of the preserved ventral domain (Extended Fig. 6C), suggesting that functionally altered niches converge on related molecular programs. Notably, the analysis of the observed expression changes between *Afadin^cKO^* and control mice for the selected gene set confirmed a significant correlation between dorsal and altered ventral regions, but not between dorsal and preserved ventral regions (Extended Data Fig. 6D).

Among shared transcriptional alterations, *Egfr*, associated with activated and dividing NSCs and TAPs^54^, was particularly prominent (Extended Data Fig. 6C, arrow). RNAscope combined with KI67 and GFAP immunostaining confirmed increased *Egfr* expression in proliferating cells in both dorsal and altered ventral V-SVZ of *Afadin^cKO^* mice (Fig. 6I, arrows). Likewise, the shared downregulation of the quiescence-associated gene *Id4* (Extended Data Fig. 6C, arrowhead) was also validated by RNAscope in both regions (Fig. 6J). qPCR analysis independently confirmed altered *Egfr* and *Id4* expression in the mutant dorsal and altered ventral V-SVZ tissue (Extended Fig. 6E).

Together, these results reveal that developmental niche disruption induces regionally distinct but partially convergent transcriptional states, with altered ventral domains acquiring molecular features associated with activated and proliferative neural stem cell programs.

### Separating intrinsic and niche-dependent effects of *Afadin* on neural stem cell activation

The increased NSC activation observed following embryonic *Afadin* deletion occurs in the context of extensive remodeling of the ventricular and ependymal architecture. We therefore asked whether *Afadin* also contributes intrinsically to NSC activation, independently of developmental alterations in the surrounding niche. To address this question, we used an *in vivo* mosaic strategy to delete *Afadin* in NSCs after birth, allowing AFADIN-deficient cells to be compared with neighboring unaffected cells within the same niche environment.

We first crossed *Afadin* floxed mice (*Afadin^F/F^*)^22^ with *Ai9* CRE-reporter mice^66^, generating homozygous double-floxed mice (*Afadin^F/F^;Ai9*) and controls lacking the floxed *Afadin* alleles (*Afadin^+/+^;Ai9*) (Fig. 7A). At P1.5, pups received intraventricular electroporation of a CAG-driven CRE plasmid to target NSCs and their progeny (Fig. 7B). In *Afadin^F/F^;Ai9* mice, CRE-mediated recombination induced permanent *tdTomato* expression and efficient *Afadin* perturbation, as confirmed by immunohistochemistry (Fig. 7C). Analysis at adult stages revealed an expansion of the *tdTomato*-labeled lineage in the V-SVZ following CRE-mediated *Afadin* deletion compared with electroporated controls (Fig. 7D-E). KI67 and GFAP immunostaining showed enhanced NSC proliferation, with a higher proportion of GFAP⁺ *tdTomato*^+^ cells co-expressing KI67 in mutants (Fig. 7F, arrows). Consistently, the OB showed a higher percentage of labeled neurons in *Afadin^F/F^;Ai9* mice (Fig. 7G-H), indicating enhanced neuronal output without an overt impairment in migration.

**Figure 7.**
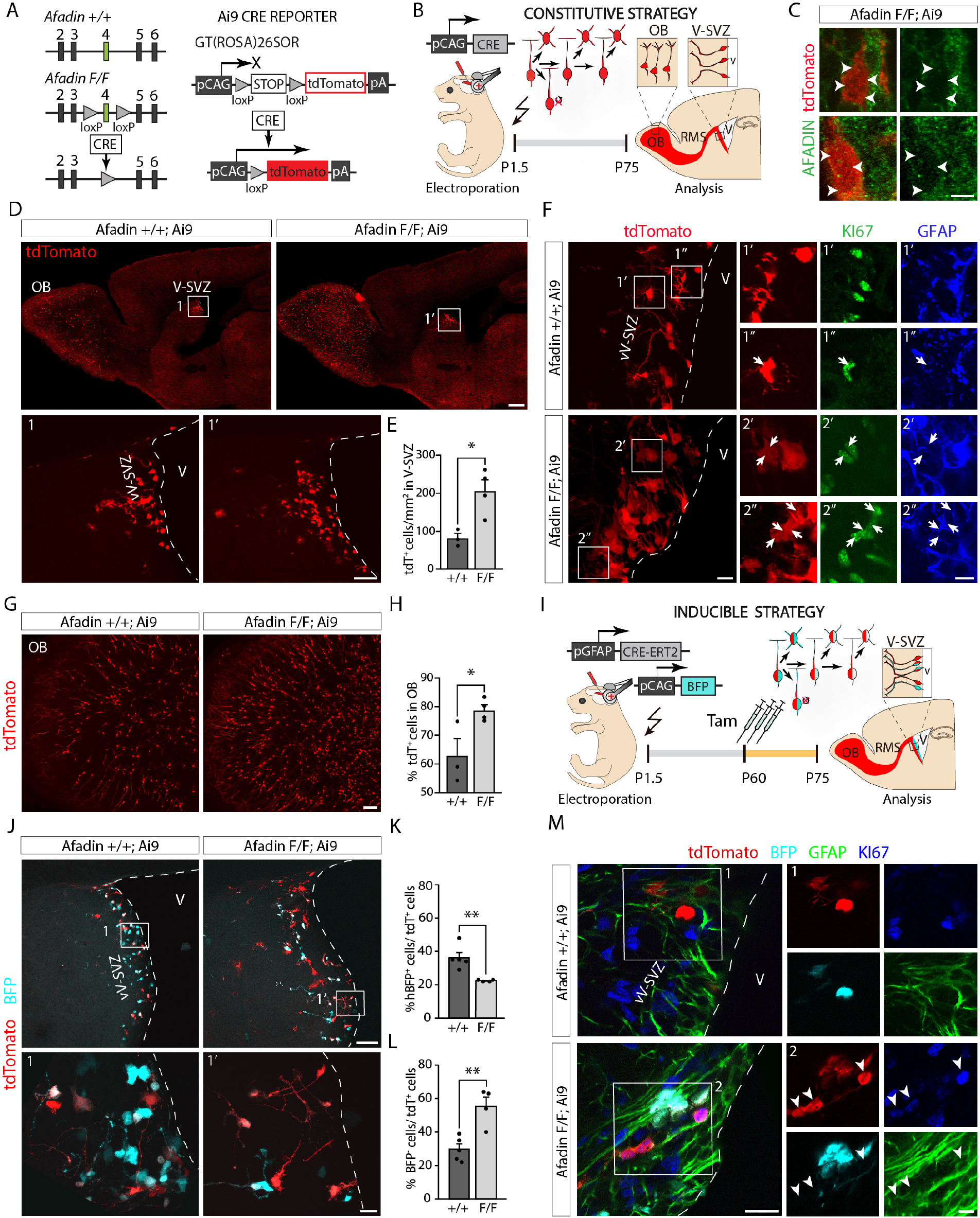
Postnatal Afadin deletion induces cell-autonomous neural stem cell activation. **A**. Schematic of the *Afadin* control (*Afadin^+/+^)*, floxed (*Afadin^F/F^*) and *Ai9*-CRE-reporter (*Ai9*) alleles. *Ai9* carries a loxP-flanked STOP cassette preventing *tdTomato* expression. **B**. Schematic of the constitutive postnatal electroporation strategy for *Afadin* inactivation. *Afadin^+/+^;* Ai9 and *Afadin^F/F^*; *Ai9* pups were electroporated with a CAG-driven CRE recombinase to permanently label NSCs and their progeny while deleting *Afadin.* **C**. *tdTomato*^+^ cells (arrowheads) in electroporated *Afadin^F/F^; Ai9* mice show loss of AFADIN expression following CRE-mediated recombination. **D-E**. Sagittal brain sections from postnatally electroporated *Afadin^+/+^; Ai9* and *Afadin^F/F^; Ai9* mice (P75) reveal an increased number of t*dTomato*^+^ cells in mutant V-SVZ (mean ± s.e.m.; *n* = 3-4; \**p* = 0.0187; unpaired two-tailed Student’s *t*-test). **F**. KI67 and GFAP immunohistochemistry showing increased NSC activation in the V-SVZ of *Afadin^F/F^; Ai9* (*tdTomato*^+^KI67^+^GFAP^+^ cells, arrows) compared with controls. **G**-**H**. Postnatal *Afadin* inactivation increases the proportion of *tdTomato*^+^ interneurons in the olfactory bulb (OB) compared with controls (mean ± s.e.m.; *n* = 3-4; \**p* = 0.0368; unpaired two-tailed Student’s *t*-test). **I**. Schematic of the inducible electroporation strategy for *Afadin* inactivation using a GFAP-driven tamoxifen-inducible CRE recombinase. Tamoxifen administration at P60 followed by a 15-day chase induces recombination in quiescent NSCs; a BFP plasmid was co-electroporated to trace cell divisions by dilution. **J**. Sagittal sections of the electroporated V-SVZ from *Afadin^+/+^; Ai9* and *Afadin^F/F^; Ai9* mice (P75) following the adult inactivation strategy show increased NSC divisions upon adult *Afadin* perturbation, visualized by a higher proportion of *tdTomato*⁺BFP⁻ cells. **K-L**. Quantification of the proportion of *tdTomato*⁺ cells with high BFP levels (hBFP, K) or lacking BFP (L) (mean ± s.e.m.; *n* = 4-5; hBFP^+^, \**p* = 0.0065; BFP^-^, \**p* = 0.0042; unpaired two-tailed Student’s *t*-test). **M**. KI67 and GFAP immunohistochemistry revealing increased NSC activation in the V-SVZ after adult *Afadin* inactivation (tdTomato^+^Ki67^+^GFAP^+^ cells, arrowheads). RMS, rostral migratory stream; vV-SVZ, ventral ventricular–subventricular zone; V, ventricle; Tam, tamoxifen. Scale bar, **C**, 5 µm; **D**, 500 µm; **D (1-1’)**, **G**, **J**, 100 µm; **F**, **J (1-1’)**, **M**, 20 µm; **F (1-2’’)**, **M (1-2)**, 10 µm.

We next asked whether *Afadin* also alter the behavior of more quiescent NSCs in adulthood. To target this population, we used a tamoxifen-inducible CRE*-*recombinase (CRE-ERT2) driven by the GFAP promoter, together with a blue fluorescent protein (BFP) reporter to monitor proliferative history. Electroporation was performed at P1.5 in *Afadin^F/F^;Ai9* and *Afadin^+/+^;Ai9*, followed by tamoxifen administration at two months of age and analysis 15 days later (Fig. 7I). Because postnatally electroporated astrocytes rapidly dilute episomal plasmids through proliferation^67,68^, this strategy preferentially labels the more quiescent NSC population that retains the construct until tamoxifen administration. BFP intensity was then used as an indirect readout of proliferative history: cells retaining high BFP levels had undergone limited proliferation, whereas BFP loss was consistent with repeated cell division (Fig. 7I). *Afadin* deletion resulted in a significant reduction in the proportion of *tdTomato^+^* cells retaining high BFP levels and a concomitant increase in *tdTomato*⁺ BFP^-^cells (Fig. 7J-L), consistent with increased cell-cycle entry and division. KI67 staining further confirmed increased proliferation among *tdTomato*^+^ GFAP^+^ cells following *Afadin* deletion (Fig. 7M, arrowheads).

Together, these experiments show that *Afadin* loss after birth is sufficient to promote NSC activation and increase neuronal output within an otherwise established niche. These findings indicate that enhanced NSC activation following *Afadin* loss can occur in the absence of widespread developmental remodeling of the niche.

## Discussion

Developmental perturbation can durably reconfigure the organization of adult neural stem cell niches. Here, we show that altering tissue organization during cortical development redirects neural progenitors into a persistent ectopic germinal zone (EGZ) while simultaneously remodeling the endogenous V-SVZ. Despite these profound anatomical changes, stem-cell activity persists into adulthood, indicating that the organization of the adult neurogenic niche is more plastic than its canonical anatomy would suggest.

The EGZ provides the clearest evidence of this plasticity. It contains NSCs, transit-amplifying progenitors, and immature neurons and supports self-renewal and multilineage differentiation into advanced adulthood. Its developmental origin is most consistent with delamination of cortical radial glial cells following disruption of adherens junction organization and apico-basal polarity ^22,47^. The accumulation of PAX6^+^ progenitors within the ectopic white matter (EWM) during the perinatal period, together with the absence of ventricular continuity, supports this interpretation. Proliferative and progenitor activity is not distributed uniformly throughout the EWM but remains concentrated in its lateralmost domain, suggesting that only a subset of displaced progenitors acquires or retains a stem-cell competence. The location of the EGZ may provide a clue to this selective persistence. It develops adjacent to the embryonic territory that contributes to the dorsal V-SVZ wedge, a specialized region of the adult niche enriched in B2 NSCs^53,69^. B2 cells are particularly relevant because they can become spatially separated from the ventricular surface while retaining long-term neurogenic potential^69^. The developmental and anatomical proximity of the EGZ to this territory raises the possibility that EGZ NSCs retain features associated with the B2 lineage or with the developmental program that gives rise to this population. Although our data do not establish such a relationship, this possibility could help explain why a subset of displaced progenitors retains long-term stem-cell properties outside the canonical ventricular niche.

The local environment may provide an additional permissive context for the persistence of stem-cell activity. The EGZ is embedded within cortical white matter and contains features associated with the canonical neurogenic niche, including vascular contacts and niche-associated microglia. Notably, the dorsal V-SVZ itself lies immediately beneath the corpus callosum, placing NSCs at the interface between the ventricular surface and a white-matter environment. The similarity in tissue context raises the possibility that the EGZ recreates, at least in part, a permissive environment for stem-cell maintenance outside its canonical anatomical position. This does not imply that anatomical location is irrelevant, but rather that stem-cell maintenance may depend on a combination of local cellular and extracellular cues that can, under some circumstances, be established outside the canonical niche. Whether vascular, glial, extracellular, or inflammatory components are sufficient to maintain the EGZ, or whether additional developmental signals are required, remains unknown.

The same developmental perturbation remodels the endogenous V-SVZ. Regions with disrupted ventricular architecture showed increased numbers of activated and proliferating progenitors, with the strongest expansion in the dorsal and adjacent ventral V-SVZ, whereas more distant ventral territories with preserved architecture showed reduced proliferative activity. This regional pattern parallels the extent of ependymal disruption: in the dorsal V-SVZ, ependymal cell specification fails, whereas in affected ventral territories ependymal cells initiate specification but fail to complete maturation. Consequently, mature multiciliated ependymal organization is not established, and GFAP^+^ cells retaining immature radial glia-like features persist along the ventricular surface. Because ependymal cells contribute to ventricular organization and regulate the NSC niche^13,70^, failure to establish this epithelial architecture may contribute to the increased stem-cell activation observed in affected domains. Consistent with this possibility, ependymal ciliary beating has recently been shown to contribute to the maintenance of adult NSC quiescence ^71^. Notably, the rostrocaudal gradient in the phenotype, with greater changes in rostral regions where cortical expansion is greater, raises the possibility that the extent of developmental tissue remodeling influences the magnitude of the adult NSC response.

Niche remodeling can also extend beyond the cells carrying the primary genetic perturbation. Although *Afadin* deletion is restricted to the dorsal lineage, adjacent ventral territories in which *Afadin* is preserved acquire altered architecture and a more activated transcriptional state. The spatial relationship between cortical expansion, ventricular deformation, and the severity of ventral remodeling is consistent with tissue-level effects, including altered epithelial coupling and mechanical constraints. Enrichment of mechanical-response and adhesion-related programs in the dorsal V-SVZ provides additional support for this interpretation. The transcriptional response follows the same regional pattern: the greatest molecular changes occur in the dorsal V-SVZ, where *Afadin* loss and architectural disruption are greatest, including reduced expression of quiescence-associated genes such as *Id4*^60^ and increased expression of genes associated with NSC activation and proliferation, including *Egfr*^54^. Ventral domains show a weaker but partially convergent response despite retaining *Afadin*, suggesting that different routes to niche remodeling can converge on related NSC states. Developmental changes in tissue organization can therefore propagate through neighboring tissue and modify stem-cell behavior beyond the primary site of genetic perturbation.

The mosaic experiments provide a complementary perspective by separating the effects of developmental niche remodeling from direct regulation of NSC state. *Afadin* deletion after birth increases NSC activation and neuronal output within an otherwise established niche, without reproducing the widespread ventricular and ependymal abnormalities caused by developmental deletion. These findings indicate that the increased NSC activity observed after developmental *Afadin* loss reflects two interacting levels of regulation: the intrinsic contribution of *Afadin* to NSC state and the broader consequences of developmental tissue remodeling. Developmental disruption therefore does not simply alter NSC behavior through changes in the surrounding niche; it also reveals an intrinsic capacity of NSCs to respond to loss of *Afadin* within an established environment. By separating the effects of intrinsic *Afadin* loss from those of developmental niche remodeling, these experiments support a model in which stem-cell state is shaped by the interaction between cell-intrinsic mechanisms and the surrounding tissue environment.

Despite sustained expansion of proliferative populations in the EGZ and remodeled V-SVZ, we did not observe overt tumor formation. Persistent activation of adult NSCs is therefore not, by itself, sufficient to drive uncontrolled growth, suggesting that additional cell-intrinsic and tissue-level constraints continue to limit stem-cell expansion^6,8^. Thus, substantial and long-lasting changes in stem-cell activity can occur without overt loss of tissue homeostasis. These changes nevertheless have functional consequences for the progeny generated by the remodeled niches. The EGZ generates neuronal progeny, some of which remain associated with the ectopic zone whereas others appear to extend beyond it, but their ultimate fate and functional integration remain unknown. In the endogenous neurogenic system, developmental *Afadin* loss disrupts RMS organization, uncoupling neuronal production from efficient migration toward the OB. In contrast, acute postnatal or adult deletion increases neuronal output without producing the same migratory defect. Impaired migration is therefore more readily explained by developmental reorganization of the surrounding tissue and migratory pathways than by loss of *Afadin* within adult NSCs themselves. Whether neurons generated in the EGZ establish stable local populations or migrate to and integrate within other brain regions remains to be determined.

The findings may also broaden the interpretation of developmental cortical malformations. The EGZ develops within the heterotopic cortical architecture of *Afadin^cKO^* mice, which exhibit a double-cortex phenotype reminiscent of human subcortical band heterotopia^26,72,73^. Such malformations have traditionally been considered primarily in terms of abnormal neuronal migration and cortical organization. Our findings suggest that the consequences of these developmental perturbations may extend beyond the initial misplacement of neurons, as early changes in tissue organization can persist into adulthood and reshape the progenitor populations that occupy the affected tissue. The persistence of a functional NSC population within this altered architecture suggests that developmental malformations may not simply displace cells but can also change the tissue context in which progenitor states are established and maintained. This observation places the EGZ within a broader conceptual framework in which the canonical anatomy of the adult niche represents a developmentally established configuration rather than an absolute requirement for stem-cell function. In this view, the long-term behavior of adult NSCs reflects both the developmental history and organization of their surrounding tissue and the intrinsic mechanisms that regulate their state.

This degree of plasticity may not be unique to neural stem-cell niches and could reflect a broader property of adult stem-cell systems. The EGZ provides a potential framework for testing whether developmental remodeling can similarly create alternative stem-cell environments in other tissues. Such principles could eventually inform strategies to establish permissive environments for endogenous or transplanted stem cells, although their potential for functional tissue regeneration will require further investigation.

Our findings therefore place developmental tissue organization as a determinant of the long-term state and configuration of adult stem-cell niches. The ability of displaced progenitors to retain stem-cell competence shows that the boundaries of the adult niche are not necessarily fixed by anatomy alone, but emerge from the interaction between developmental history, tissue organization, and intrinsic cellular programs.

## Materials and Methods

### Mice

All animal procedures were conducted in accordance with Spanish legislation, European Union Directive 2010/63/EU, and U.S. animal welfare regulations. Experimental protocols were approved by the Ethical Committee of the University of Valencia and the *Conselleria de Agricultura, Desarrollo Rural, Emergencia Climática y Transición Ecológica* (Comunidad Valenciana).

*Afadin^F/F^* mice (*Afadin^Ctl^*) were previously described by Gil-Sanz *et al*. (2014) and kindly provided by Dr. Ulrich Müller. Conditional knockout mice *(Afadin^cKO^*) were generated by crossing *Afadin^F/F^* animals with *Emx1*-Cre mice^74^, originally obtained from The Jackson Laboratory (005628), and kindly provided by Dr. Ulrich Müller. To generate *Afadin^F/F^;Ai9* and *Afadin^+/+^;Ai9* mice, *Afadin^F/F^* animals were crossed with *Ai9^F/F^* Cre-reporter mice^66^ (Jackson Laboratory, 007909), also kindly provided by Dr. Ulrich Müller.

All mouse lines were maintained on a C57BL/6J background and bred at the University of Valencia’s animal facility (Burjassot). Both male and female mice were used in all the experiments of this work. Genotyping was performed by PCR using standard protocols.

### *In vivo* Manipulations

#### Thymidine analog injections

Adult *Afadin^Ctl^* and *Afadin^cKO^* mice (P80) received seven intraperitoneal (i.p.) injections of 5-ethynyl-2’-deoxyuridine (EdU; Santa Cruz, sc-284628A) at 50 mg/kg body weight, administered at 2-hour intervals. Mice were sacrificed 30 days later to identify cells undergoing limited divisions or label-retaining cells (LRCs)^19,39,75^.

#### *In vivo* postnatal electroporation

P1.5 *Afadin^+/+^;Ai9* and *Afadin^F/F^;Ai9* pups were anesthetized by hypothermia^75^. A plasmid solution (1.5 µg/µl DNA) in endotoxin-free TE buffer (QIAGEN, 1018499) with 10% Fast Green (Sigma-Aldrich, F7252-5G) was injected into one lateral ventricle. Electroporation was performed using the ECM830 Square Wave electroporator (BTX, 45-0052) with 7 mm platinum tweezer electrodes (BTX, 45-0488), delivering five electric pulses (90 V, 90 ms duration, 950 ms interval). Pups were recovered on a heating pad and returned to the dam. For inducible electroporation experiments, P60 electroporated mice received three i.p. injections of tamoxifen (Sigma-Aldrich, T5648-1G) at 75 mg/kg body weight, administered at 12-hour intervals to induce CRE recombination. In both paradigms, mice were analyzed at P75. DNA constructs pCAG-CRE, pCAG-CRE-ERT2 and pCAG-BFP were donated by the laboratory of Dr. Ulrich Müller. pGFAP-CRE-ERT2 was subcloned in our laboratory.

#### FlashTag injection

P0.5 *Afadin^Ctl^* and *Afadin^cKO^* pups were anesthetized by hypothermia, as described above, and injected into a lateral ventricle with 1 µl of a solution containing 1/10 volume of 10 mM Green *FlashTag* or carboxylic acid diacetate succinimidyl ester (CellTrace™ Oregon Green™ 488; Thermo Fisher, C34555) diluted in endotoxin-free TE buffer supplemented with 10% Fast Green. Pups were recovered on a heating pad, returned to the dam, and sacrificed 1 h later for analysis.

### Tissue Processing and Histology

#### Tissue processing

Postnatal and adult mice were deeply anesthetized and transcardially perfused with 4% paraformaldehyde (PFA) in 1X phosphate-buffered saline (PBS), except for P0 pups, whose brains were dissected and immersion-fixed overnight at 4 °C. Brains were post-fixed, stored in PBS with 0.05% sodium azide at 4 °C, and sectioned either on a vibratome (Leica, VT1200S; 40 µm standard sections; 1 mm *en face* preparations) after embedding in 4% low-melting-point agarose, or on a cryostat (Leica, CM1900) at 16 µm following cryoprotection in 30% sucrose and embedding in O.C.T. (CellPath, KMA-0100-00A) for RNAscope. For whole-mount preparations, the V-SVZ of non-perfused adult mice was dissected and immersion-fixed overnight in 4% PFA at 4 °C without agitation.

#### Immunohistochemistry

Sections were incubated for 1 h at room temperature (RT) in blocking solution (10% horse serum, 0.2% Triton X-100 in 1X PBS), followed by primary antibodies overnight at 4 °C. After washing in PBS, sections were incubated with secondary antibodies and DAPI (1:1000) for 1 h at RT and mounted with Fluoromount-G (Electron Microscopy Sciences, 17984-25). EdU detection was performed using the Click-iT^TM^ EdU Alexa Fluor^TM^ 555 Imaging Kit (ThermoFisher, C10338). For *en face* and whole-mount preparations, tissues were blocked for 4 h and incubated with primary antibodies for 48 h at 4 °C without agitation, followed by secondary antibodies for 2h at RT. *En face* sections were microdissected under a microscope to isolate dorsal and ventral V-SVZ regions. *En face* and whole-mount preparations were mounted to expose the ventricular surface with Fluoromount G. Antibodies and dilutions are listed in **Supplementary Table 1-2**.

#### RNAScope *in situ* hybridization

Cryostat brain sections were processed according to the manufacturer’s instructions for the RNAscope™ Multiplex Fluorescent Reagent Kit v2 (Advanced Cell Diagnostics, Bio-Techne, 323100). To combine RNAscope *in situ* hybridization with immunohistochemistry, sections were washed with 1X PBS after the final RNAscope conjugation step and subsequently processed for immunohistochemistry as described above. Gene specific probes used: *Egfr* (Bio-techne, 1579451-C1) and *Id4* (Bio-techne, 447861-C2).

### Imaging and Quantification

Images were acquired with a FluoView FV10i confocal laser-scanning microscope (Olympus) and LSM 980 super-resolution confocal microscope (Zeiss). Image processing and quantitative analyses were performed using Adobe Photoshop and Fiji (ImageJ), except for 3D V-SVZ reconstructions, which were generated using Arivis, and lateral ventricle reconstructions, which were generated using ITK-SNAP (v.4.0.0). For quantification, 3–4 sections per animal were analyzed across different rostrocaudal levels. In *Afadin^cKO^*V-SVZ, altered ventral and preserved ventral regions were defined based on differences in GFAP immunoreactivity patterns.

For 3D reconstruction and volumetric analysis of the lateral ventricles, MRI images were imported into ITK-SNAP and ventricular boundaries were manually delineated in all coronal and axial sections from each animal. The software automatically calculated ventricular volumes from the segmented regions and generated the corresponding three-dimensional reconstructions^76^.

For RMS analysis, DCX-stained consecutive sagittal sections were reconstructed in Photoshop to generate complete RMS projections, then binarized and analyzed for area and positive pixel count using Fiji. Orientation of migratory chains was assessed using the OrientationJ plugin (Fiji, Cubic Spline method, α = 2°), generating color-coded maps and histograms^77,78^.

For OB analyses, reconstructed full sections were analyzed in Fiji. OB layers were delineated on DAPI images to measure areas and dimensions. DCX^+^ signal was binarized to quantify positive pixels, and EdU^+^ LRCs and *tdTomato*^+^ interneurons were detected using the “Find Maxima” function after thresholding to exclude background.

### Ultrastructural and MRI Analyses

#### Scanning Electron Microscopy (SEM)

Dorsal and ventral V-SVZ *en face* preparations were post-fixed in osmium tetroxide, dehydrated through an ethanol gradient (70–100%), and dried via critical point drying (Tousimis, Autosamdri-814). Samples were mounted on metal stubs and sputter-coated with gold-palladium (1 min, 1500 V, 10–20 mA) using a Quorum/Polaron SC7640 system. Imaging was performed on a Hitachi S4800 Scanning electron microscope (SCSIE, University of Valencia) at 1 kV using 3000× magnification.

#### Magnetic Resonance Imaging (MRI)

MRI scans were acquired on a Bruker Avance III 400 WB spectrometer (400 MHz, 9.4 T) equipped with a wide-bore microimaging probe (SCSIE, University of Valencia, U26 NMR: Biomedical Applications II platform from Nanbiosis, Research Infrastructures & Services of CIBER-BBN). Perfused mouse heads were imaged in 2.5 cm glass tubes. After low-resolution T1-weighted scans for positioning, Turbo RARE T2-weighted sequences were collected across 25 axial, coronal, and sagittal planes (field of view, 20×20 mm; matrix, 512×512; slice thickness, 250 µm), yielding a final resolution of 39×39×250 µm. T2-weighted Turbo RARE 3D sequences were acquired and coronal plane images (field of view, 12×12 mm; matrix, 512×512; slice thickness, 100 µm) were reconstructed from 3D sequences with Simple Slices (MPR) task in ParaVision (v.6.0.1, Bruker). Temperature was kept at 15 °C during the MRI experiments.

### Transcriptomic Analyses

#### RNA extraction

Dorsal, upper-ventral and lower-ventral V-SVZ regions, as well as EGZ and corresponding control subregions were microdissected from 1-mm-thick coronal vibratome sections, at intermediate positions along this axis, of *Afadin^Ctl^*and *Afadin^cKO^* brains under a dissecting microscope. Total RNA was extracted using the RNeasy Plus Micro Kit (Qiagen, 74034). mRNA was reverse transcribed and preamplified using the SuperScript IV Single Cell/Low Input cDNA PreAmp Kit (Invitrogen, 11752048) following manufacturer’s guidelines, and cDNA was purified using AMPure XP beads (Beckman Coulter, A63880).

#### RNA-seq

Libraries were prepared and sequenced by Novogene on an Illumina NovaSeq X platform (paired-end, 150 bp). All libraries were sequenced to a sequencing depth of >= ≥ 20M read pairs per sample. Reads were aligned with hisat2 (v2.0.5)^79^ to Ensembl release 94 reference genome. Differential expression analysis was performed with DESeq2 package ^80^. Gene ontology analysis was performed using R package clusterProfiler^81^.

#### qPCR

Gene expression was quantified by real-time quantitative PCR (qPCR) using a StepOnePlus™ Real-Time PCR System (Applied Biosystems). Reactions were performed using either gene-specific primer pairs with TB Green Premix Ex Taq™ (Takara, RR420A) or TaqMan™ probes with TaqMan™ Fast Advanced Master Mix (Applied Biosystems, 4444557). Reference genes were selected based on stable expression across experimental conditions. For TB Green assays, *Atp5f1* was used as the reference gene, whereas *Gapdh* was used for TaqMan assays. For both approaches, 4–10 ng of cDNA were used in a final reaction volume of 10 μL, and amplification was performed using a standard 40-cycle program with annealing and extension at 60 °C. Primer sequences are listed in **Supplementary Table 3**.

### Neurosphere Culture and *In Vitro* Assays

#### Neurosphere culture

Adult (P60) and aged (P240) *Afadin^Ctl^* and *Afadin^cKO^* mice were sacrificed by cervical dislocation. Brains were embedded in 4% low-melting-point agarose (Fisher BioReagents, BP16525) and coronal sections (1 mm) were cut with a vibratome for microdissection of the EGZ and an equivalent cortical region. Tissue was enzymatically dissociated for 30 min at 37 °C in papain solution (12 U/mouse; Worthington, LS003119) containing 0.2 mg/ml L-cysteine hydrochloride (Sigma-Aldrich, C8277), and 0.2 mg/ml EDTA (Sigma-Aldrich, E6511) in EBSS (Gibco, 24010-043), and carefully triturated with a Pasteur pipette to obtain a single-cell suspension. Cells were collected by centrifugation and plated at 15,000 cells/cm² in DMEM/F12 medium (Gibco, 11320-074) modified as previously described and supplemented with 20 ng/ml epidermal growth factor (EGF; Gibco, 53003-018) and 10 ng/ml fibroblast growth factor 2 (FGF2; Sigma-Aldrich, F0291) ^36,37^. Cultures were maintained at 37 °C with 5% CO₂ for 7 DIV, after which primary neurospheres were counted and imaged using an inverted microscope (Nikon Eclipse TE2000-S).

### In vitro assays

For culture expansion, neurospheres were dissociated with Accutase® (Sigma-Aldrich, A6964) and replated in the same medium (EGF and FGF2) at 10,000 cells/cm². Cultures were passaged every 5–6 DIV for up to 10 passages. For proliferation and differentiation assays under adherent conditions, 40,000 dissociated cells/cm² were seeded on Matrigel-coated (Corning, 354234), pre-flamed coverslips in the same medium without EGF. After 2 DIV, FGF2 was removed and cells were cultured for an additional 5 days in medium without mitogens, supplemented with 2% heat-inactivated fetal bovine serum (HI-FBS; Biowest, S181B-500) to induce differentiation^37^. Cultures were fixed with 2% PFA for 15 min at both 2 and 5 DIV and stored in PBS with 0.05% sodium azide.

### Immunocytochemistry and imaging

Coverslip-attached cells were incubated for 30 min at RT in blocking solution containing 0.1% Triton X-100, followed by incubation with primary antibodies (3 h) and secondary antibodies (1 h), and counterstaining with DAPI (1:1000). For O4 detection, a triple-layer immunostaining protocol was used without Triton X-100, using biotinylated goat anti-mouse IgG (Jackson ImmunoResearch, 115-066-072; 1:1000) followed by Cy2-streptavidin (Jackson ImmunoResearch, 016-220-084; 1:2000). Coverslips were mounted using Fluoromount G, and images were acquired with a FluoView FV10i confocal microscope. For quantification, 25-30 randomly selected fields of view from 3-4 different coverslips per animal were analyzed using Photoshop to determine the percentage of different cell populations. Antibodies and dilutions are listed in **Supplementary Table 1-2**.

### Statistical Analysis

All statistical analyses were performed using GraphPad Prism (v9.5.0). The statistical test used for each experiment is indicated in the corresponding figure legend. Normality was tested using the Shapiro–Wilk test. Parametric data were compared with the Student’s *t*-test, and non-parametric data were analyzed with the Mann–Whitney test. Experiments involving two independent variables and repeated measurements were analyzed using two-way repeated-measures ANOVA followed by Šídák’s multiple-comparisons test. Statistical significance was set at *p* < 0.05. Significance in figures is indicated as follows: \**p* < 0.05, \*\**p* < 0.01, \*\*\**p* < 0.001, \*\*\*\**p* < 0.0001, and exact *p* values are reported in the figure legends. Data are presented as mean ± s.e.m., with individual data points shown. Sample sizes (*n* = 3–6) are indicated in each figure legend.

## Acknowledgments

We thank Dr. Isabel Fariñas and Dr. Isabel Martínez-Garay for access to reagents and equipment. We acknowledge Dr. Isabel Martinez-Garay and Santos Franco for critical review of the manuscript. We thank the Servicio Central de Soporte a la Investigación Experimental (SCSIE-UVEG), especially the animal production, microscopy, and NMR sections. U26 NMR: Biomedical Applications II platform from Nanbiosis (Research Infrastructures & Services of CIBERBBN) are also gratefully acknowledged.

This work was supported by grants from the Ministerio de Ciencia e Innovación (PID2020-114227RB-I00, MICIU/AEI/10.13039/501100011033; CNS2022-135758, MICIU/AEI/10.13039/501100011033 and the European Union NextGenerationEU/PRTR; and PID2023-153143OB-I00, MICIU/AEI/10.13039/501100011033 and FEDER, EU) awarded to C.G.-S., and by grants PID2022-142734OB-I00 (MICIU/AEI/10.13039/501100011033 and FEDER, EU) and EUR2023-143479 (MICIU/AEI/10.13039/501100011033) awarded to S.R.F. Additional funding was provided by the Generalitat Valenciana (CIAICO/2024/287 to C.G.-S. and S.R.F.). I.M.-W. was the recipient of a Garantía Juvenil contract from the Generalitat Valenciana (GJIDI/2018/A/221). L.L.-C. (PRE2020-094137) was the recipient of a predoctoral contract from the Ministerio de Ciencia e Innovación. A.M.-G. (CIACIF/2021/187), L.V.-E. (CIACIF/2024/209), C.M.M.-M. (CIACIF/2022/366), and J.F.-B. (ACIF/2021/139) are or were recipients of predoctoral contracts from the Generalitat Valenciana.

## Author Contribution

Conceptualization, I.M-W., C.G-S.; Methodology, I.M-W., C.G-S., A. M-G., L.V-E., J.F-B., L.L-C., J.P., CM.M-M., MC.M-B., SR.F.; Investigation, I.M-W., C.G-S., A. M-G., L.V-E., J.F-B., L.L-C., J.P., CM.M-M., MC.M-B., E.M-M., SR.F. Writing Original Draft, C.G-S.; Writing – Review & Editing, I.M-W., C.G-S., A. M-G., L.V-E., J.F-B., L.L-C., J.P., CM.M-M., MC.M-B., E.M-M., SR.F.; Supervision, C.G-S.; Project Administration, C.G-S.; Funding Acquisition: C.G-S., SR.F.

## Competing interests

The authors declare no competing interests.

## Extended Data Figure Legends

**Extended Data Fig. 1.**
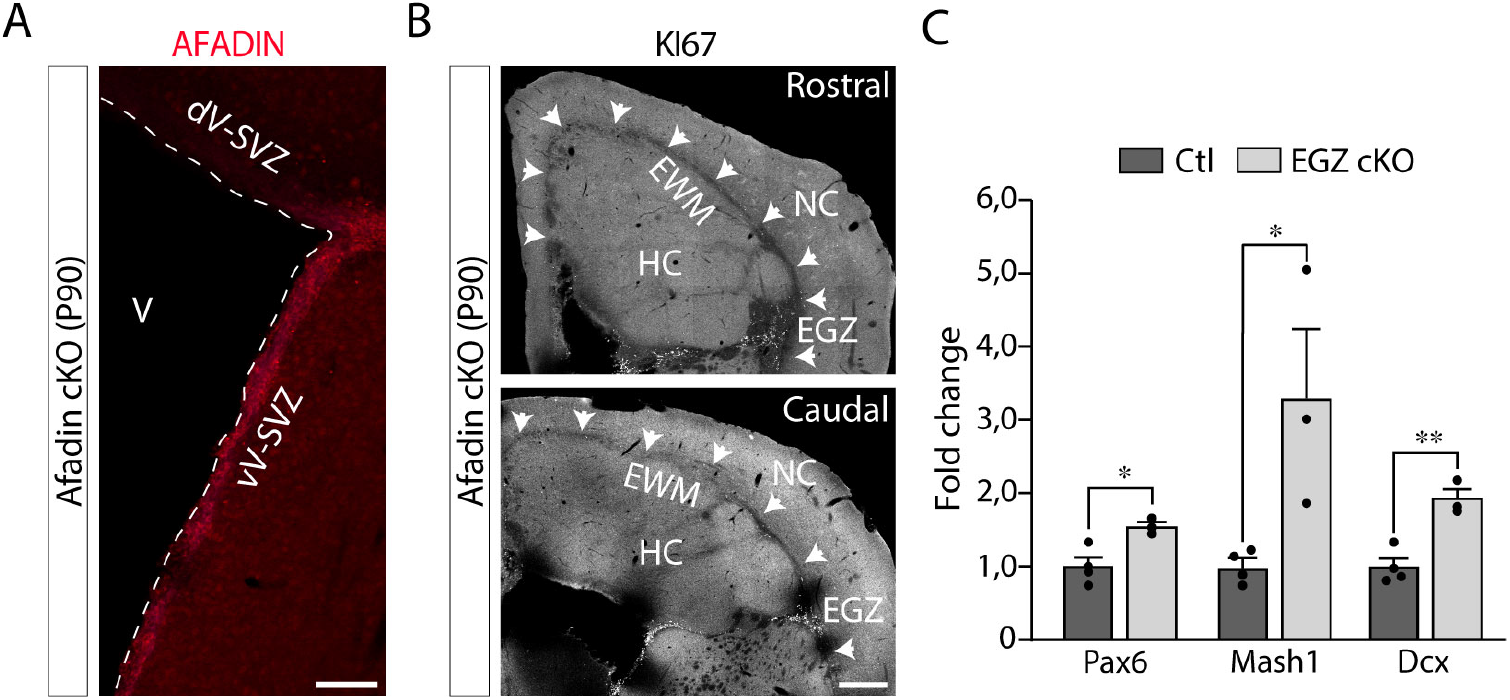
Regional AFADIN expression, spatial restriction, and neurogenic signature of the adult EGZ. **A**. AFADIN expression is lost in the dorsal V-SVZ but preserved in ventral regions of *Afadin^cKO^* mice (P90). **B**. Across the rostrocaudal axis of adult *Afadin^cKO^* mice, KI67⁺ cells within the ectopic white matter (EWM, arrowheads) are confined to its lateralmost region, which defines the ectopic germinal zone (EGZ). **C**. qPCR analysis of neurogenic markers in the adult EGZ compared with wild-type cortical tissue (mean ± s.e.m.; *n* = 3–4; *Pax6*, \**p* = 0.017; *Ascl1*, \**p* = 0.033; *Dcx*, \*\**p* = 0.0035; unpaired two-tailed Student’s *t*-test). HC, heterocortex; NC, normocortex; Str, striatum; d/vV-SVZ, dorsal/ventral ventricular– subventricular zone; V, ventricle. Scale bars: **A**, 100 µm; **B**, 500 µm.

**Extended Data Fig. 2.**
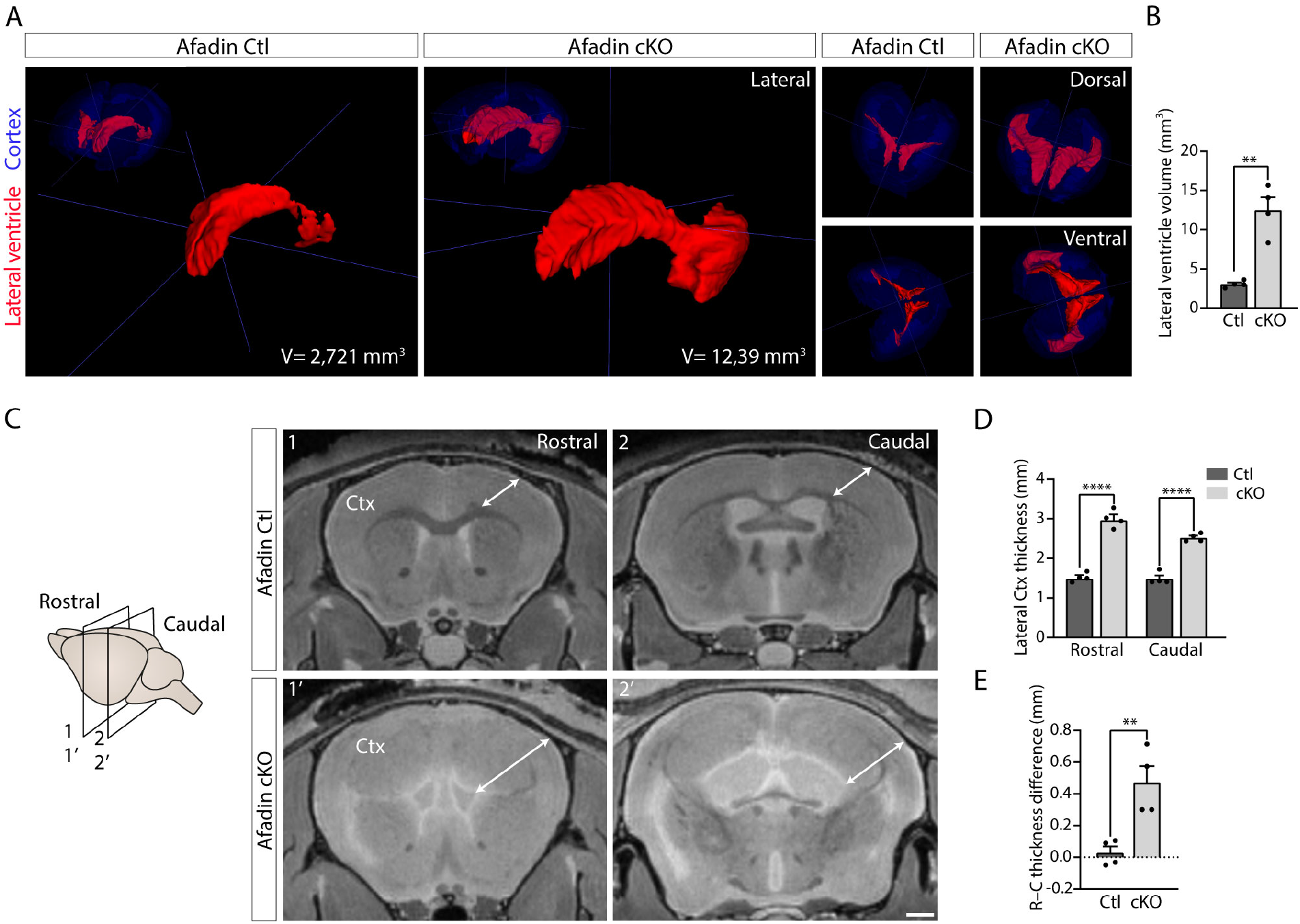
Regional changes in ventricular architecture accompany V-SVZ niche remodeling. **A**. Three-dimensional reconstruction of adult control and *Afadin^cKO^*cortices (blue) generated using *ITK-SNAP* software, highlighting the lateral ventricles (red) in lateral (1, 1’, 2, 2’), dorsal (3, 3’) and ventral (4, 4’) views. Note the differences in ventricular shape and size in mutant mice. **B**. Quantification of the lateral ventricle volume in control and *Afadin^cKO^*mice (mean ± s.e.m.; *n* =4; \*\**p* < 0.001; unpaired two-tailed Student’s *t*-test). **C**. Analysis of differential cortical expansion along the rostrocaudal axis in *Afadin^cKO^* mice using MRI. **D**. Quantification of lateral cortical thickness from the data in C (mean ± s.e.m.; *n* =4; Rostral, \*\*\*\**p* < 0.0001; Caudal, \*\*\*\**p* < 0.0001; two-way repeated-measures ANOVA with Šídák’s multiple-comparisons test). **E**. Quantification of the difference between rostral and caudal cortical thickness in control and *Afadin^cKO^* mice from the data shown in C (mean ± s.e.m.; *n* =4; \*\**p* = 0.0071; unpaired two-tailed Student’s *t*-test). Ctx, cortex; V, volume. Scale bars: **A (1-1’**,**3-3’**, **4-4’)**, 2mm; **A (2-2’)**, **C**, 1 mm. MRI voxel size: 0.07812 × 0.07812 × 0.09375 mm (X × Y × Z), defining the spatial resolution of the 3D reconstructions. All distances shown are in real physical space derived directly from voxel dimensions (no rescaling).

**Extended Data Fig. 3.**
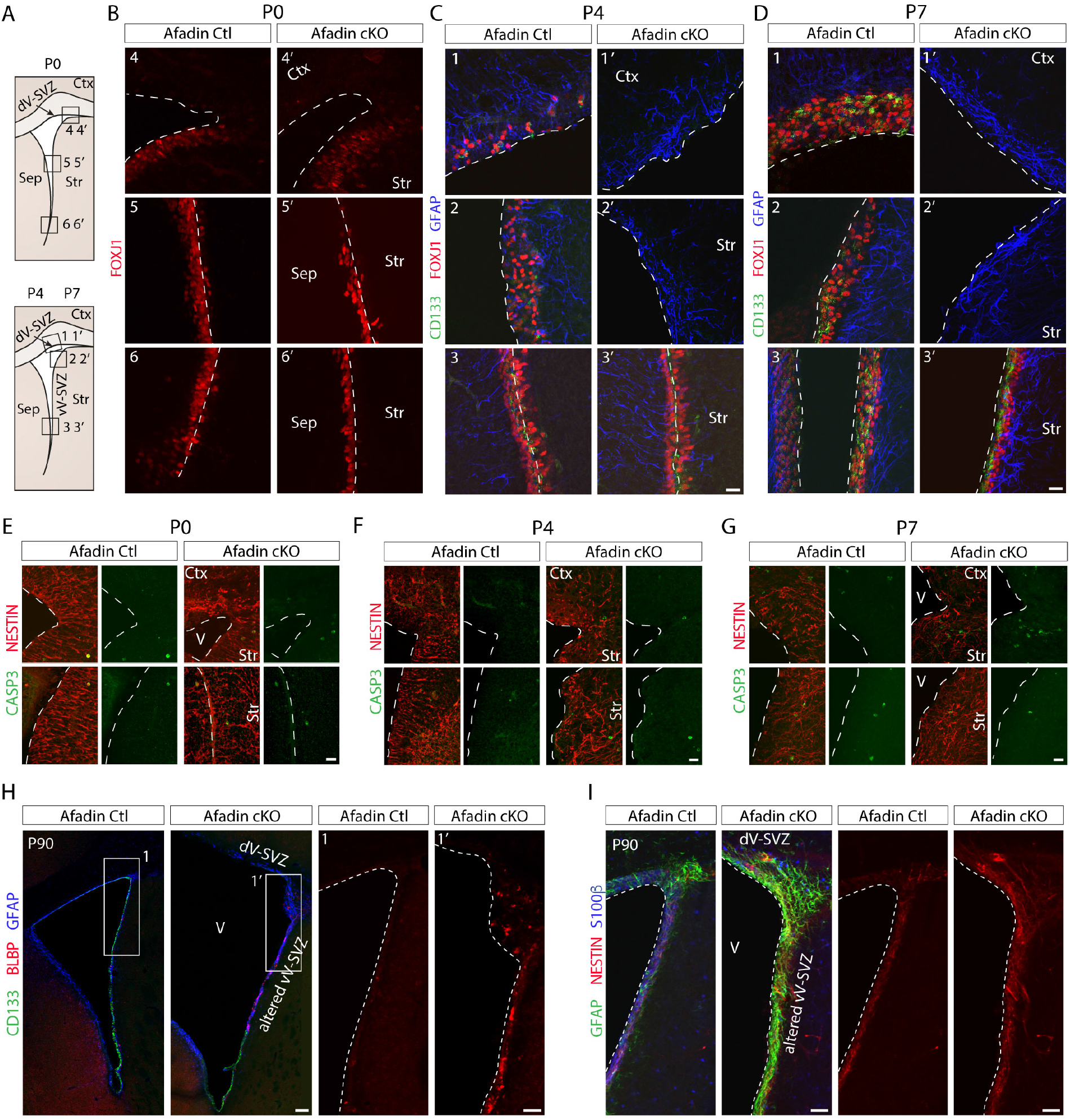
Regional differences in ependymal specification and differentiation drive V-SVZ niche reorganization. **A.** Schematic of the V-SVZ subregions analyzed in coronal sections from control and *Afadin^cKO^* mice at postnatal stages P0, P4, and P7. **B**. At P0, the onset of FOXJ1 expression is detected in the upper ventral V-SVZ of both control and *Afadin^cKO^* mice. Signal intensity was enhanced in the upper ventral V-SVZ for visualization of early FOXJ1 expression. **C-D**. Immunohistochemistry for FOXJ1, CD133 and GFAP at P4 (B) and P7 (C) reveals loss of ependymal cell markers and an increase in GFAP^+^ cells in the dorsal and the altered ventral V-SVZ of mutants. **E-G**. Immunohistochemistry for cleaved CASPASE-3 (CASP3) and the NSC marker NESTIN in the dorsal and altered ventral V-SVZ shows no major differences in the abundance of CASP3*^+^*cells between control and *Afadin^cKO^* mice at P0 (E), P4 (F) and P7 (G). **H**. BLBP immunohistochemistry in the adult V-SVZ (P90) reveals increased BLBP expression in *Afadin^cKO^* regions lacking ependymal markers, identified by GFAP^+^CD133^-^immunostaining, compared with controls. **I**. NESTIN expression is elevated in the dorsal and altered ventral V-SVZ of *Afadin^cKO^* mice, identified by GFAP^+^S100β^-^immunostaining. Ctx, cortex; d/vV-SVZ, dorsal/ventral ventricular–subventricular zone; Sep, septum; Str, striatum; V, ventricle. Scale bars: **A-F**, 20 µm; **G**, 100 µm; **G (1-1’)**, **H**, 50 µm.

**Extended Data Fig. 4.**
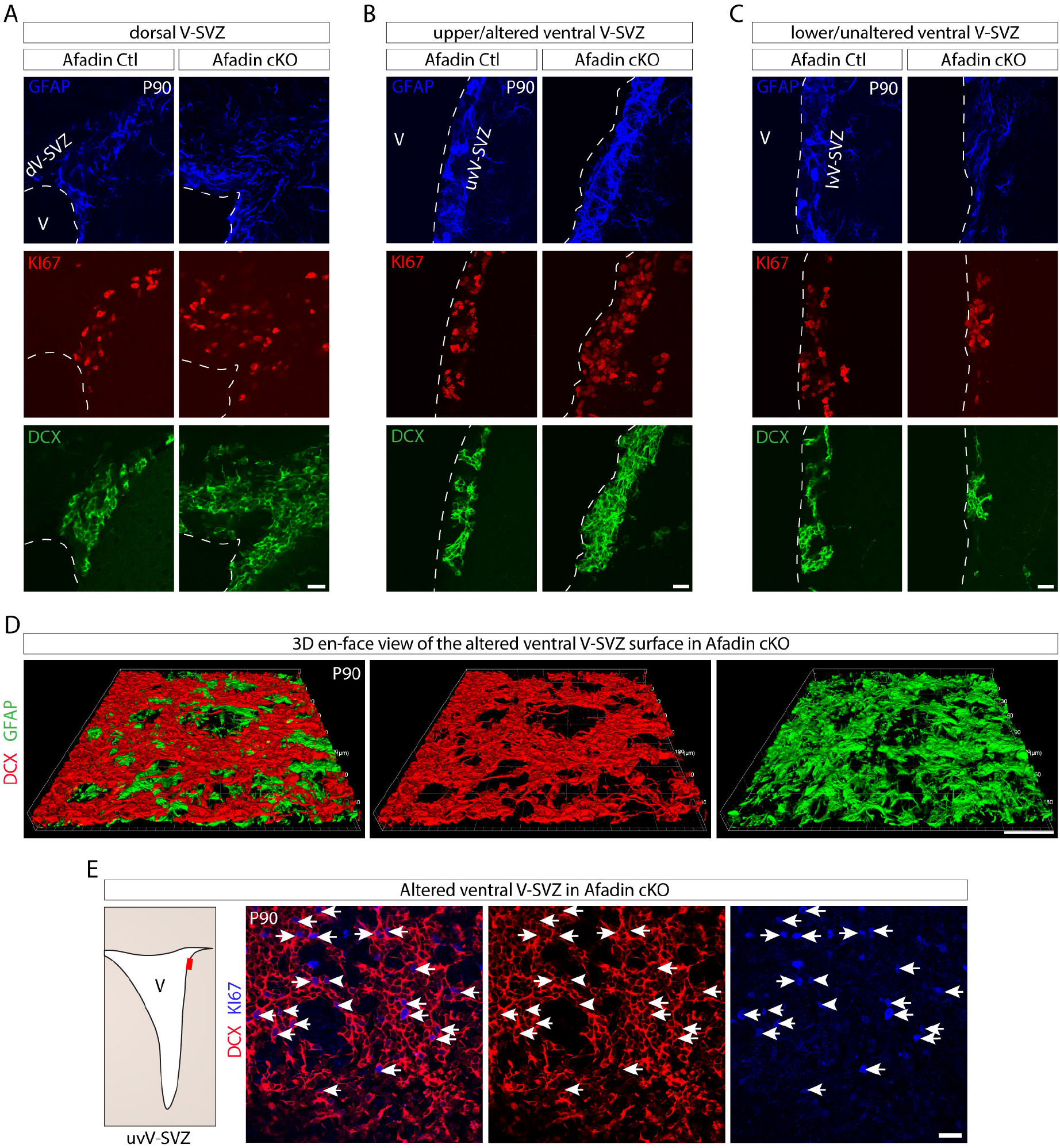
Changes in V-SVZ organization and neurogenic cell distribution. **A-C**. Immunohistochemical analysis of GFAP, KI67 and DCX in dorsal (A), altered ventral (B) and preserved ventral (C) regions of the V-SVZ from *Afadin^Ctl^* and *Afadin^cKO^*mice (P90), revealing region-specific alterations in neurogenic population in *Afadin^cKO^* mice. These panels correspond to the magnified regions shown in Fig. 5B, G, L, and display single-channel representations for clarity. **D**. Three-dimensional reconstruction of the altered ventral V-SVZ using *Arivis* software confirms accumulation of DCX⁺ cells at the ventricular surface, positioned above GFAP⁺ cells in *Afadin^cKO^*mice. **E**. *En face* analysis shows that a substantial fraction of DCX⁺ cells located at the ventricular surface co-express the proliferation marker KI67 (arrows). V, ventricle; d/ vV-SVZ, dorsal/ventral ventricular–subventricular zone. Scale bars: **A-C**, **E**, 20 µm; **D**, 30 µm.

**Extended Data Fig. 5.**
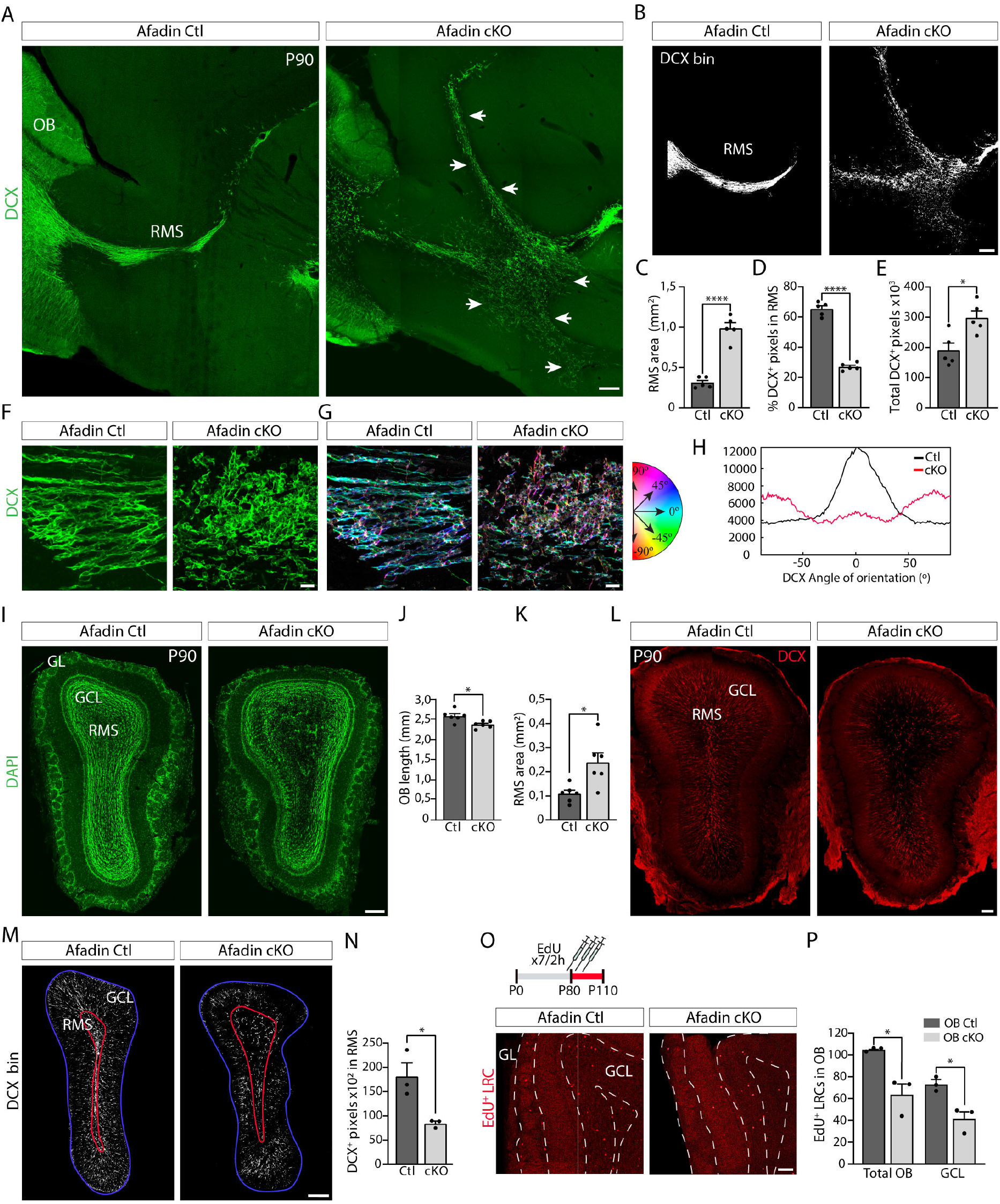
Altered V-SVZ organization impairs RMS integrity and neuronal incorporation into the olfactory bulb. **A**. DCX immunohistochemistry in sagittal sections reveals altered organization of DCX^+^ cells along the rostral migratory stream (RMS) in *Afadin^cKO^*mice, including the presence of alternative migratory routes (arrows) absent in controls. **B**. Binarized reconstructions of the RMS generated from DCX immunostaining of serial sections from the same animal. **C-E**. Quantification of the binarized reconstructions shows increased RMS area (C), reduced DCX^+^ cell density (D), and increased total DCX^+^ signal (E) in *Afadin^cKO^* mice, compared with controls (mean ± s.e.m.; *n* = 5; RMS area, \*\*\*\**p* < 0.0001; % DCX^+^ pixels in RMS, \*\*\*\**p* < 0.0001; Total DCX^+^ pixels, \**p* = 0.0116; unpaired two-tailed Student’s *t*-test). **F**. Higher magnification of the RMS highlights disorganized DCX^+^ chains in mutant mice. **G**-**H**. Orientation analysis of DCX^+^ chains using *ImageJ* plugin *OrientationJ*. Color-coded maps (G) and corresponding histograms (H) show aligned orientations in controls and a broader distribution in *Afadin^cKO^* mice. **I-K**. DAPI staining reveals overall preservation of the olfactory bulb (OB) architecture, despite changes in OB length (J) and RMS area (K) in mutant mice (mean ± s.e.m.; *n* = 6; OB length, \**p* = 0.0118; RMS area, \**p* = 0.0115; unpaired two-tailed Student’s *t*-test). **L**-**N**. DCX immunohistochemistry and binarized image quantification indicate impaired arrival of immature neurons to the OB in *Afadin^cKO^* mice (mean ± s.e.m.; n = 3; DCX^+^ pixels in RMS, \**p* = 0.0249; unpaired two-tailed Student’s *t*-test). **O-P**. EdU pulse–chase analysis reveals reduced incorporation of newborn interneurons into the *Afadin^cKO^* OB, particularly in the granular cell layer (GCL), compared with controls (mean ± s.e.m.; n = 3; Total OB, \**p* = 0.0146; GCL, \**p* = 0.0151; unpaired two-tailed Student’s *t*-test). GL, glomerular layer; Scale bars: **A**, **B**, **I**, **L**, **M**, 200 µm; **F**, **G**, 20 µm; **O**, 50 µm.

**Extended Data Fig. 6.**
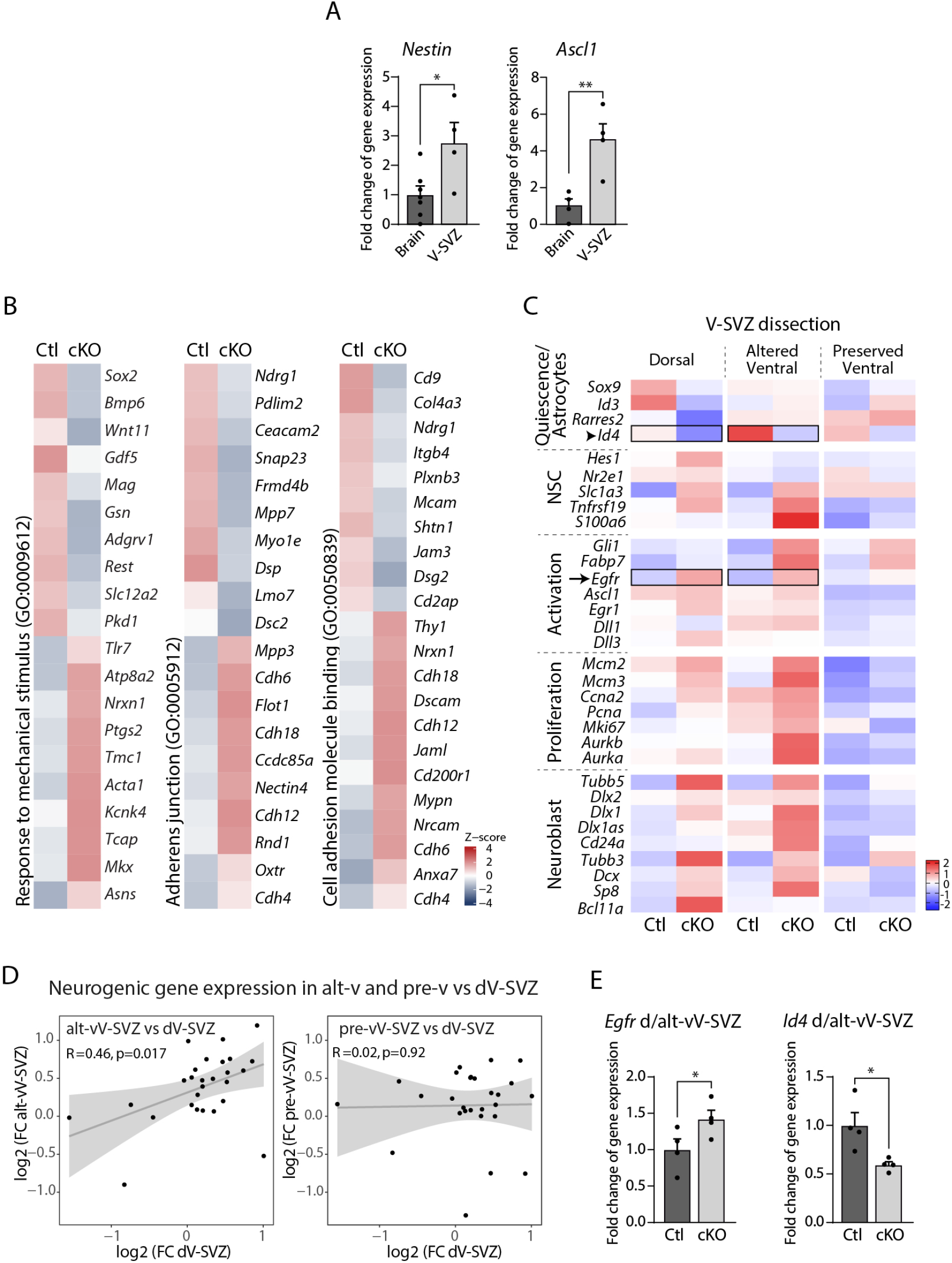
Validation and extended characterization of remodeled V-SVZ transcriptional states. **A.** qPCR validation of microdissected V-SVZ tissue of control mice shows enrichment of neurogenic markers (*Nestin, Ascl1*) relative to whole brain, confirming accurate niche isolation (mean ± s.e.m.; n = 4-7; *Nestin* \**p* = 0.0236; *Ascl1*, \*\**p* = 0.009; unpaired two-tailed Student’s *t*-test). **B.** Heatmaps of Z-score–scaled expression of DEGs in dorsal V-SVZ for selected biological processes identified by GO analysis, separated into upregulated and downregulated transcripts in *Afadin^cKO^* versus control mice. **C**. Heatmaps of DEGs associated with defined V-SVZ subpopulations ^55,65^ in control and *Afadin^cKO^* mice across V-SVZ subregions, highlighting *Egfr* (arrow) and *Id4* (arrowhead). **D.** Spearman correlation analysis based on the selected gene sets in D shows significant similarity between dorsal and altered ventral V-SVZ, but not between dorsal and preserved ventral regions. *R* and *p* values are indicated. **E.** qPCR validation confirms upregulation of *Egfr* and downregulation of *Id4* in the dorsal and altered ventral *Afadin^cKO^* V-SVZ compared to controls (mean ± s.e.m.; n = 3–4; *Egfr*, \**p* = 0.0357; *Id4*, \**p* = 0.0119; unpaired one-tailed Student’s *t*-test). d/alt/pre-vV-SVZ, dorsal/altered ventral/preserved ventral ventricular-subventricular zone.

## Notes

### Competing Interest Statement

The authors have declared no competing interest.

### Summary of Updates

This version of the manuscript has been revised to further clarify the conceptual implications of our findings. We have revised the text to better highlight stem cell plasticity, including the capacity of stem cells to generate new niches at ectopic sites and to reorganize the endogenous niche. We have also expanded the discussion to consider how these principles of niche remodeling and stem cell plasticity could potentially apply to other stem cell systems. In addition, we have reorganized and adjusted several figures to improve the presentation and clarity of the data. Some figures and panels have been moved to the Supplementary Information to focus the main figures on the findings most relevant to the central message of the study. Overall, these revisions aim to strengthen the conceptual framework and broader implications of our findings.

